# The biocide triclosan induces (p)ppGpp dependent antibiotic tolerance and alters SarA dependent biofilm structures in *Staphylococcus aureus*

**DOI:** 10.1101/2023.02.01.525840

**Authors:** Dean Walsh, Andrea Salzer, Christiane Wolz, Jonathan Aylott, Kim R Hardie

**Affiliations:** School of Life Sciences, University of Warwick, Coventry, UK; Interfaculty Institute of Microbiology and Infection Medicine, University of Tübingen, Tübingen, Germany; Cluster of Excellence EXC 2124 “Controlling Microbes to Fight Infections”, University of Tübingen; Boots Science Building, University of Nottingham, Nottingham, UK; Biodiscovery Institute, University of Nottingham, Nottingham, UK

**Keywords:** Biofilms, Antibiotic resistance, Antibiotic tolerance, Biocides, Triclosan, *Staphylococcus aureus*, stringent response, SarA, polysaccharide intercellular adhesin, SigB

## Abstract

The biocide triclosan is used extensively in both household and hospital settings. The chronic exposure to the biocide occurring in individuals that use triclosan-containing products results in low levels of triclosan present in the human body that has been linked to induction of antibiotic tolerance and altered biofilm formation. Here we aimed to unravel the molecular mechanisms involved in triclosan induced antibiotic tolerance and biofilm formation in *Staphylococcus aureus*. Triclosan treatment prior to planktonic exposure to bactericidal antibiotics resulted in 1,000 fold higher viable cell counts compared to non-pretreated cultures. Triclosan pretreatment also protected *S. aureus* biofilms against otherwise lethal doses of antibiotics as shown by live/dead cell staining and viable cell counting. Triclosan mediated antibiotic tolerance in planktonic and biofilm cultures required an active stringent response because a pppGpp^0^ strain was not protected from antibiotic killing. Incubation of *S. aureus* with triclosan also altered biofilm structure due to SarA-mediated overproduction of the polysaccharide intercellular adhesin (PIA) in the biofilm matrix. Thus, physiologically relevant concentrations of triclosan can trigger (p)ppGpp dependent antibiotic tolerance as well as SarA dependent biofilm formation.

**Importance:** The prevalent bacterium *Staphylococcus aureus* infects skin lesions and indwelling devices, and this can cause sepsis with 33% mortality. Intrinsic to this is the formation of co-ordinated communities (biofilms) protected by a polysaccharide coat. *S. aureus* is increasingly difficult to eradicate due to its antibiotic resistance. Protection against Methicillin Resistant *S. aureus* (MRSA) includes pre-hospital admission washing with products containing biocides. The biocide triclosan is the predominant antibacterial compound in sewage in Ontario due to its use in household and hospital settings. Levels of triclosan accumulate with exposure in humans. The significance of our research is in identifying the mechanisms triggered by exposure of *S. aureus* to physiological levels of triclosan that go on to raise the tolerance of *S. aureus* to antibiotics and promote the formation of biofilms. This understanding will inform future criteria used to determine effective antimicrobial treatments.

## Introduction

Biofilms are surface attached communities of bacteria enclosed in an exopolymeric matrix. For the pathogen *Staphylococcus aureus*, this matrix is composed of surface proteins (1), polysaccharide intercellular adhesin (PIA) (2), and extracellular DNA (eDNA) (3, 4). Biofilms can act as reservoirs of antibiotic tolerance and use numerous mechanisms to withstand antimicrobial treatment, including reduced penetration of antimicrobials into the biofilm matrix (5), and a generally slow growth rate (6). This reduced rate of growth serves to protect bacteria by diminishing the efficacy of the majority of antimicrobials. The enduring resilience of biofilms has made them a severe clinical concern, particular with regards to chronic infections caused by biofilm forming bacteria such as *S. aureus* (7).

Antibiotic tolerance is the process by which an entire bacterial population can survive transient exposure to antibiotics that would otherwise be lethal (8, 9). Antibiotic tolerance distinguishes itself from antibiotic resistance in numerous ways. For example, resistant bacteria may need higher concentrations of an antimicrobial to achieve bacterial killing, whereas tolerant bacteria require a longer exposure time to an antimicrobial to provide the same level of killing achieved against susceptible bacteria (8). Additionally, resistant bacteria are typically protected against a single antibiotic or a small group of closely related antibiotics, whilst tolerant bacteria can typically better withstand a broad array of antimicrobials (8, 9). The presence of tolerant bacterial populations in clinical settings results in the widespread misclassification of these bacteria as resistant (8), leading to flawed antimicrobial treatments and risking recurrent infection in patients (10, 11). Antibiotic tolerance in *S. aureus* is becoming increasingly relevant clinically, as exemplified by 6%-43% of *S. aureus* clinical strains being vancomycin tolerant (12). Drug tolerant *S. aureus* can cause prolonged fevers and extended bacteraemia duration, increased treatment failures and even increased mortality rates (13, 14).

There is a broad range of molecular strategies employed by *S. aureus* to facilitate antibiotic tolerance (15). The stringent response is one such strategy and is conserved among most bacterial species as a means to combat nutritional deficiencies (16, 17). During the stringent response the alarmones ppGpp and pppGpp (collectively known as (p)ppGpp) are synthesised from ATP and GTP (pppGpp) or GDP (ppGpp). In *S. aureus*, alarmone synthesis is driven by three alarmone synthetases: two small alarmone synthetases; RelP and RelQ, and a larger protein Rel. Rel is bifunctional and equipped with a synthetase domain for alarmone synthesis and a hydrolase domain for the purpose of alarmone degradation. Hydrolysis of these alarmones in *S. aureus* is essential for survival, as accumulation of (p)ppGpp results in cell death (18, 19). Rel, RelP, and RelQ are triggered by different inducing stresses, including various antibiotics. Numerous studies have used the antibiotic mupirocin, an antibiotic that induces amino acid starvation, to induce Rel activity (18, 20, 21). RelP and RelQ expression has been triggered by exposure to the cell wall targeting antibiotics ampicillin, oxacillin, and vancomycin (20, 22). Once the stringent response has been triggered, (p)ppGpp orchestrates global changes in gene expression, with steep downregulation of genes associated with proliferative functions, such as protein synthesis and DNA replication (23, 24). As bactericidal antibiotics disrupt the function of active targets, the metabolic shutdown and quiescence induced by the stringent response renders these drugs largely ineffectual (25).

The synthetic biocide triclosan inhibits type II fatty acid synthesis (FASII) at the enoyl-acyl carrier protein reductase (FabI) step (26, 27). At high concentrations, triclosan provokes membrane permeability and non-specific membrane damage (28). In hospitals, triclosan is used as an MRSA decolonisation therapy (29), antimicrobial hand wash (29), antiseptic ointment (30), and is impregnated into surgical sutures (31) and urinary catheters (32). Triclosan is also widely used in domestic settings, being found in many household products including soaps, toothpastes (33), cosmetics and laundry detergents (34). This widespread use of triclosan means that both people and environments are chronically exposed to the biocide. Absorption of triclosan from triclosan containing toothpastes (TCT) is high, with triclosan blood concentration increasing from 0.81 ng/mL to 296 ng/mL after 14 days of TCT use (35). The average urine concentration of triclosan in patients exposed to the biocide during their hospital stay was 245 ng/mL, with concentrations reaching as high as 505 ng/mL (33).

Recent work suggests that pretreatment with low levels of triclosan can induce antibiotic tolerance in both *Escherichia coli* and *S. aureus* to multiple different antibiotics (36). Moreover, insufficiently dosed treatments can alter biofilm formation (37). *S. aureus* has been shown to grow thicker biofilms in the presence of sub-MIC mupirocin (38), clindamycin (39), and β-lactams (40) due to increased eDNA release, whilst vancomycin exposure caused increased biofilm formation due to elevated levels of both PIA and eDNA in the biofilm matrix (41). Salicylic acid, the active component of aspirin, has been shown to increase the production of PIA in *S. aureus* biofilms (42). Although triclosan has not yet been observed to alter the biofilm formation of *S. aureus*, the biocide does stimulate cellulose production in established *Salmonella typhimirium* biofilms (43). As biocides are globally accessible and both biofilm formation and antibiotic tolerance have been associated with antibiotic treatment failure, augmentation or induction of these processes by biocide exposure could be a serious clinical concern.

This study aimed to investigate how triclosan exposure affects not just planktonic *S. aureus*, but also *S. aureus* biofilms. We show that low levels of triclosan can induce antibiotic tolerance via the stringent response in *S. aureus* biofilms, thereby increasing their resilience against antibiotic killing. In addition, triclosan exposure throughout biofilm formation led to a significant increase in biofilm polysaccharide production in a SarA-dependent manner. Overall, low levels of triclosan, consistent with those found accumulating within the human body, can trigger multiple molecular mechanisms to drastically alter the phenotype of *S. aureus* biofilms, protecting them against antibiotics and possibly additional threats.

## Materials and Methods

### Bacterial strains and culture conditions

*S. aureus* was grown at 37°C on brain heart infusion (BHI) agar plates for 16-20 hours. For growth in liquid media *S. aureus* was grown at 37°C in BHI broth with shaking at 200 rpm unless stated otherwise.

**Table 1.** S. aureus strains used in this study

| Strain | Description | Reference |
| --- | --- | --- |
| HG001 | Derivative of NCTC8325;<br>*rsbU* repaired, *tcaR*<br>defective, carries<br>prophages $\Phi 11$, $\Phi 12$, and<br>$\Phi 13$. | (44) |
| HG001 (p)ppGpp<sup>0</sup> | HG001 rel/relP/relQ triple<br>mutant ($\Delta rel_{syn}$, $\Delta relP_{syn}$,<br>$\Delta relQ_{syn}$) | (20) |
| HG001 *sarA* | HG001 *sarA*::<br>*Bursa aurealis*<br>(*erm*) | (23) |
| NCTC8325-4 | Derivative of NCTC8325;<br>*rsbU* and *tcaR* defective,<br>cured of prophages $\Phi 11$,<br>$\Phi 12$, and $\Phi 13$. | (45) |
| HG001 $\Delta sigB$ | HG001<br>$\Delta mazEFrsbUVWsigB::tetM$ | (46) |

### Antibiotic susceptibility testing

#### Antibiotics tested

Antimicrobials used for these experiments were obtained from Sigma-Aldrich: triclosan (Igrasan), ciprofloxacin, vancomycin, rifampicin.

#### Minimum inhibitory concentration (MIC) determination

96 well microtitre plates were inoculated with *S. aureus* strains at an optical density of 600 nm (OD_600_) of 0.05 in Mueller-Hinton broth. Minimum inhibitory concentrations (MICs) were determined by the broth microdilution method in accordance with EUCAST guidelines (47).

#### Minimum biofilm eradication concentration (MBEC) determination

MBEC experiments were conducted by inoculating 96 well microtitre plates with *S. aureus* at an OD_600_ of 0.05 in Mueller-Hinton broth and incubating for 24 hours at 37°C to allow biofilm formation. Following this, a 2-fold dilution series of antibiotics was added to wells and biofilms incubated in Mueller-Hinton broth with antibiotics for a further 24 hours at 37°C. After incubation, biofilms were removed from wells by vigorous scraping with pipette tips and sonicating at 30 kHz for 1 minute in sonication bath. Disrupted biofilms were then spotted out onto Mueller-Hinton agar plates and the MBEC defined as the minimum antibiotic concentration at which viable colonies are not detected.

### Time-kill assays

*S. aureus* strains at OD_600_ 0.01 were inoculated into either untreated BHI broth or BHI broth supplemented with triclosan (500 ng/mL; 0.25x MIC). For conditions requiring fatty acid supplementation, 500 μM oleic acid and 0.1% v/v Brij 58 for solubilisation of oleic acid was also added. Cultures were then incubated at 37°C in shaking conditions (200 rpm) for a period of 30 minutes to allow for acclimatisation to biocide and/or fatty acid supplementation.

Following this, inhibitory concentrations of antibiotics were added to relevant conditions. Throughout this study the antibiotics ciprofloxacin (1 μg/mL), vancomycin (2 μg/mL), rifampicin (40 ng/mL) were used. This resulted in four conditions per strain being present in each experiment, consisting of untreated *S. aureus*, biocide exposed, antibiotic treated, biocide exposed + antibiotic treated. Cultures were then returned to the incubator and samples were taken every hour for 6-10 hours depending on the experiment. Each hour, colony forming units (CFUs) were determined and the OD_600_ recorded to determine viability and growth, respectively.

### Biofilm imaging

#### Biofilm preparation

Overnight cultures of *S. aureus* were diluted in BHI to OD_600_ 0.05 and loaded into μ-slide 8-well glass bottomed chambers (ibidi, glass bottom). 500 ng/mL triclosan, was added to relevant wells, allowing biofilms to form in the presence of triclosan. For conditions requiring fatty acid supplementation, 500 μM oleic acid and 0.1% Brij 58 was also added. Chambers were then placed in a 37°C incubator for 48 hours in static conditions. Following this incubation period, growth medium was removed from the biofilms and replaced with fresh BHI growth medium.

#### Live/dead staining

To investigate antibiotic tolerance either 4096 μg/mL ciprofloxacin, 2048 μg/mL vancomycin or 2048 μg/mL rifampicin were added to wells for 3 hours. Next, live/dead staining was carried out using 6 μM syto 9 and 30 μM propidium iodide (BacLight, Molecular Probes). Biofilms were imaged using a Zeiss LSM 700 compact confocal laser scanning microscope using the 40x objective and appropriate fluorescence settings (syto 9=488 nm laser, propidium iodide=555 nm laser).

#### Staining of biofilm components

To examine biofilm structure, the nucleic acid stain 4’,6-diamidino-2-phenylindole (DAPI) was added at a final concentration of 10 μg/mL and the polysaccharide stain fluorescein-conjugated wheat germ agglutinin (WGA) (Invitrogen™) was added at a final concentration of 10 μg/mL. Biofilms were imaged using a Zeiss LSM 700 compact confocal laser scanning microscope using the 40x objective and appropriate fluorescence settings (DAPI=405 nm laser, fluorescein=488 nm laser).

#### Quantification using Comstat2 software

Biomass of live cells, dead cells, and stained matrix polysaccharide was analysed using Comstat2 software (48).

### Biofilm characterisation

#### Biofilm preparation

0.5 mL of BHI was added to 24 well plates and inoculated with OD_600_ 0.05 *S. aureus*. Wells were treated with either 500 ng/mL triclosan, 500 μM oleic acid solubilised in 0.1% Brij 58, or a combination of both triclosan and solubilised oleic acid. Biofilms were grown statically at 37°C for 48 hours. For biofilm experiments including antibiotics, 4096 μg/mL ciprofloxacin, 2048 μg/mL vancomycin or 2048 μg/mL rifampicin were added to wells for the last 12 hours of biofilm incubation. Following incubation, growth medium was removed and biofilms washed with PBS.

#### Crystal violet staining to quantify biomass

Following initial incubation the protocol was carried out as described previously (49) with minor alterations. Briefly, 200 μL of 0.1% crystal violet was added to biofilms for 15 minutes. Crystal violet was then removed and biofilms washed with PBS. Biofilms were then left to dry for 1 hour and then crystal violet was solubilised using 30% acetic acid. Absorbance values at 550 nm were then measured.

#### Calcofluor white staining to quantify polysaccharide

Calcofluor white staining was carried out as previously described (50) with minor alterations. 200 μL of calcofluor white (1 mg/mL in dH_2_O) was added to wells and incubated in the dark for 1 hour. Calcofluor white was then removed and biofilms washed with PBS to remove unbound calcofluor white. 200 μL of 96% ethanol was then added to solubilise biofilm-bound calcofluor white. Fluorescence intensity was then measured using the 360 nm excitation filter and 460 nm emission filter.

#### CFU determination

PBS was added to wells and biofilms were sonicated at 30 kHz for 1 minute, diluted, and CFUs determined. For experiments involving antibiotics, the % viability of the biofilm was defined as the CFU of the antibiotic treated sample (for instance WT+cip or WT T+cip) divided by the CFU of the corresponding control (WT untreated or WT T). This corrected for inherent differences in cell number when comparing an untreated *S. aureus* biofilm to a triclosan exposed *S. aureus* biofilm.

## Results

### Triclosan induces antibiotic tolerance towards ciprofloxacin and vancomycin, but not rifampicin in planktonically grown *S. aureus*

Triclosan was added to planktonic *S. aureus* cultures at a concentration of 500 ng/mL (5× MIC HG001; Table S1), corresponding to physiologically relevant concentrations of triclosan found in the urine of users of triclosan-containing products (51). Triclosan pretreatment provided protection from inhibitory concentrations of ciprofloxacin (4× MIC) (Fig 1B) and vancomycin (Fig 1D). Viable cell counts in triclosan-exposed (T+C, T+V) conditions remained relatively constant throughout the time-kill assays, whilst conditions treated with ciprofloxacin (C) or vancomycin (V) alone displayed a sharp decrease in cell viability over time. Triclosan-induced protection resulted in a 100-fold increase in cell survival in the presence of ciprofloxacin, a DNA gyrase inhibitor, by 3 hours of antibiotic exposure, and a 1000-fold difference by 5 hours (T+C, Fig 1B). Likewise, triclosan exposure also provided 1000-fold higher cell survival in the presence of the cell wall synthesis targeting antibiotic vancomycin (1× MIC) by 6 hours (T+V, Fig 1D). The growth profiles in Fig 1A and 1C show that all treatments (T, C, V, and combinations) lead to drastic inhibition of growth, despite triclosan being administered at a physiologically relevant dose. The mismatch between optical density and log_10_CFU/mL should be noted. For instance, although the T, C, T+C conditions prevented detectable broth growth, biocide pretreated bacteria exhibited 2-3 log higher CFUs than ciprofloxacin only treated bacteria. Notably, triclosan was unable to induce antibiotic tolerance against rifampicin (4× MIC) (Fig S1B), despite the combination of triclosan and rifampicin displaying the same pattern of growth inhibition seen in previous experiments with ciprofloxacin or vancomycin (Fig S1A). This may suggest that triclosan-induced tolerance only protects against bactericidal antibiotics such as ciprofloxacin and vancomycin, but not bacteriostatic antibiotics such as rifampicin.

**Figure 1.**
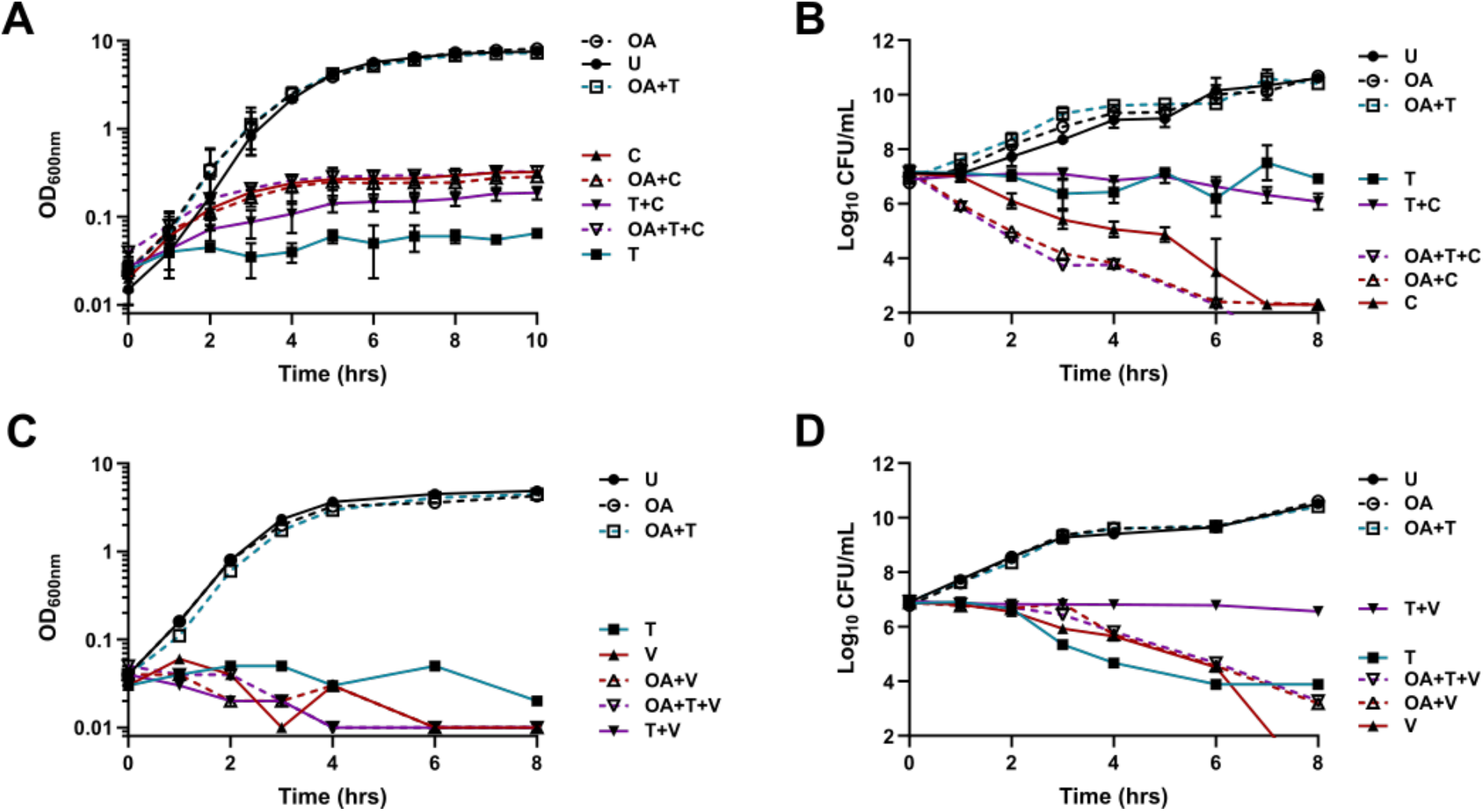
Pretreatment with triclosan protects *S. aureus* planktonic cultures from antibiotic treatment, this protection is negated by oleic acid supplementation. For planktonic experiments, *S. aureus* strain HG001 was incubated in BHI in the presence or absence of triclosan, with or without 500 μM oleic acid solubilised in 0.1% Brij 58, for 30 mins at 37°C. Inhibitory concentrations of antibiotics were added and incubation continued. Untreated (U, OA), 500 ng/mL triclosan pretreated (T, OA+T), 1 μg/mL ciprofloxacin treated (C, OA+C), 500 ng/mL triclosan treated and 1 μg/mL ciprofloxacin treated (T+C, OA+T+C), 2 μg/mL vancomycin treated (V, OA+V), 500 ng/mL triclosan treated and 2 μg/mL vancomycin treated (T+V, OA+T+V) conditions are shown. Growth curves with ciprofloxacin **(A)** and vancomycin **(C)** showing optical densities (OD_600_) plotted against time. Error bars display ±SD, n=3. **B)** Ciprofloxacin time-kill assays, **D)** vancomycin time-kill assays with Log_10_ CFUs plotted against time. Error bars display ±SD, n=3.

Antibiotic tolerance is often associated with nutritional starvation (52). As triclosan is a fatty acid synthesis inhibitor, it was investigated whether oleic acid supplementation could counter fatty acid starvation caused by triclosan, thereby preventing triclosan-induced antibiotic tolerance. A physiologically relevant concentration of oleic acid (500 μM), consistent with the levels of fatty acids found in human serum (53), was added alongside triclosan pretreatment. The data presented in Fig 1B and Fig 1D suggest that fatty acid starvation caused by triclosan plays a role in protecting the bacteria against ciprofloxacin and vancomycin. This is evidenced by oleic acid supplementation causing triclosan pretreated cultures to be just as susceptible to ciprofloxacin (OA+T+C) and vancomycin (OA+T+V) as cultures that received no triclosan pretreatment (OA+C, OA+V). Oleic acid had no effect on the growth or viability of untreated or antibiotic alone conditions (OA, OA+C, OA+V, OA+R) (Fig 1A, 1C, S1A). OA+T conditions demonstrated growth and viability similar to an untreated sample, demonstrating that the presence of fatty acids negates the growth inhibition caused by physiologically relevant concentrations of triclosan.

### Triclosan can protect *S. aureus* biofilms from eradication by high concentrations of ciprofloxacin, vancomycin, and rifampicin

Following the findings that triclosan exposure could induce antibiotic tolerance in planktonic *S. aureus*, it was investigated whether triclosan exposure could induce antibiotic tolerance in *S. aureus* biofilms also. Biofilms grown in the presence of triclosan and treated with ciprofloxacin (1× MBEC) (T+C, Fig 2A), vancomycin (1× MBEC) (T+V, Fig S2B) and rifampicin (1× MBEC) (T+R, Fig S2C) displayed significantly less cell death compared to biofilms that were not pretreated with triclosan (C, V, R) (Fig 2B). Just as in planktonic experiments, oleic acid supplementation alongside triclosan pretreatment prevented triclosan-induced antibiotic tolerance, with OA+T+C, OA+T+V, and OA+T+R biofilms having live cell percentages comparable to C, V, and R (Fig 2B). Live dead microscopy shows that triclosan alone (T; Fig 2A), oleic acid alone (OA; Fig S2A) and the fatty acid with triclosan (OA+T; Fig S2A) had little to no effect on cell viability compared to untreated biofilms (U) (Fig 2B).

**Figure 2.**
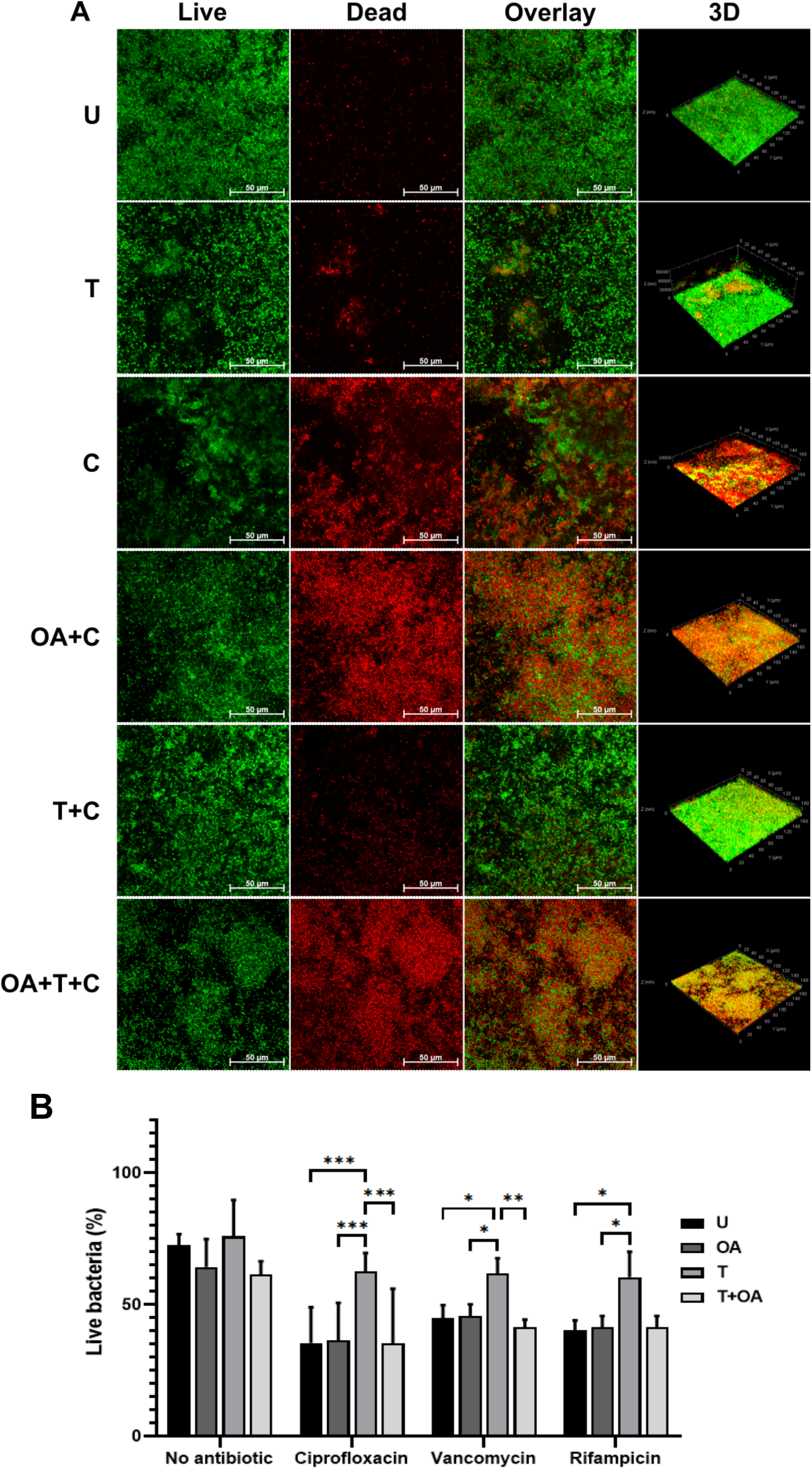
Triclosan exposure protects *S. aureus* biofilms from antibiotic treatment, this protection is negated by oleic acid supplementation. CLSM was used to assess the effect of triclosan on the antibiotic tolerance of *S. aureus* HG001 WT. Syto 9 (green) was used to visualise live cells, whilst propidium iodide (red) was used to visualise cell death. **A)** Untreated biofilms (U), 500 ng/mL triclosan exposed biofilms (T), biofilms grown in the presence or absence of 500 μM oleic acid and treated with 4096 μg/mL ciprofloxacin (C, OA+C) and 500 ng/mL triclosan exposed biofilms treated with 4096 μg/mL ciprofloxacin (T+C, OA+T+C) are shown. For each condition a 2D image of a selected z-plane is shown for live, dead, and overlay images. A 3D image of each condition is also shown. Images are representative of multiple experiments and were taken using the 40x objective (n = 3). **B)** Quantification of the percentage of live cells in biofilms was carried out using Comstat2 image analysis software. Error bars represent SD, n=3. *** denotes P≤0.001, ** denotes P≤0.01, * denotes P≤0.05.

### Exposure to triclosan throughout biofilm formation results in increased polysaccharide production

Fluorescent staining was used to characterise the composition of triclosan exposed *S. aureus* biofilms (Fig 3A). Alongside DAPI to visualise cells, fluorescein-conjugated WGA was used to stain the PNAG residues of the PIA polysaccharide that is prevalent in the *S. aureus* biofilm matrix. CLSM revealed that triclosan exposure significantly alters the production of polysaccharide in the *S. aureus* biofilm matrix. Echoing the findings of previous antibiotic tolerance experiments, oleic acid supplementation was able to largely negate the triclosan-induced changes, with enhanced polysaccharide production not being observed in OA+T biofilms. Comstat2 quantification showed that triclosan biofilms are composed of significantly more WGA stained PNAG compared to untreated (U), oleic acid only (OA) or oleic acid and triclosan (OA+T) pretreatments (Fig 3B), indicative of more PIA polysaccharide. Despite triclosan exposed biofilms producing more PIA, overall biomass as determined by crystal violet staining was significantly lower than unexposed biofilms (Fig S3B, S3C). This is because triclosan exposure resulted in a significantly lower population of cells present (Fig S3A) due to the reduced growth rate noted in Fig 1A.

**Figure 3.**
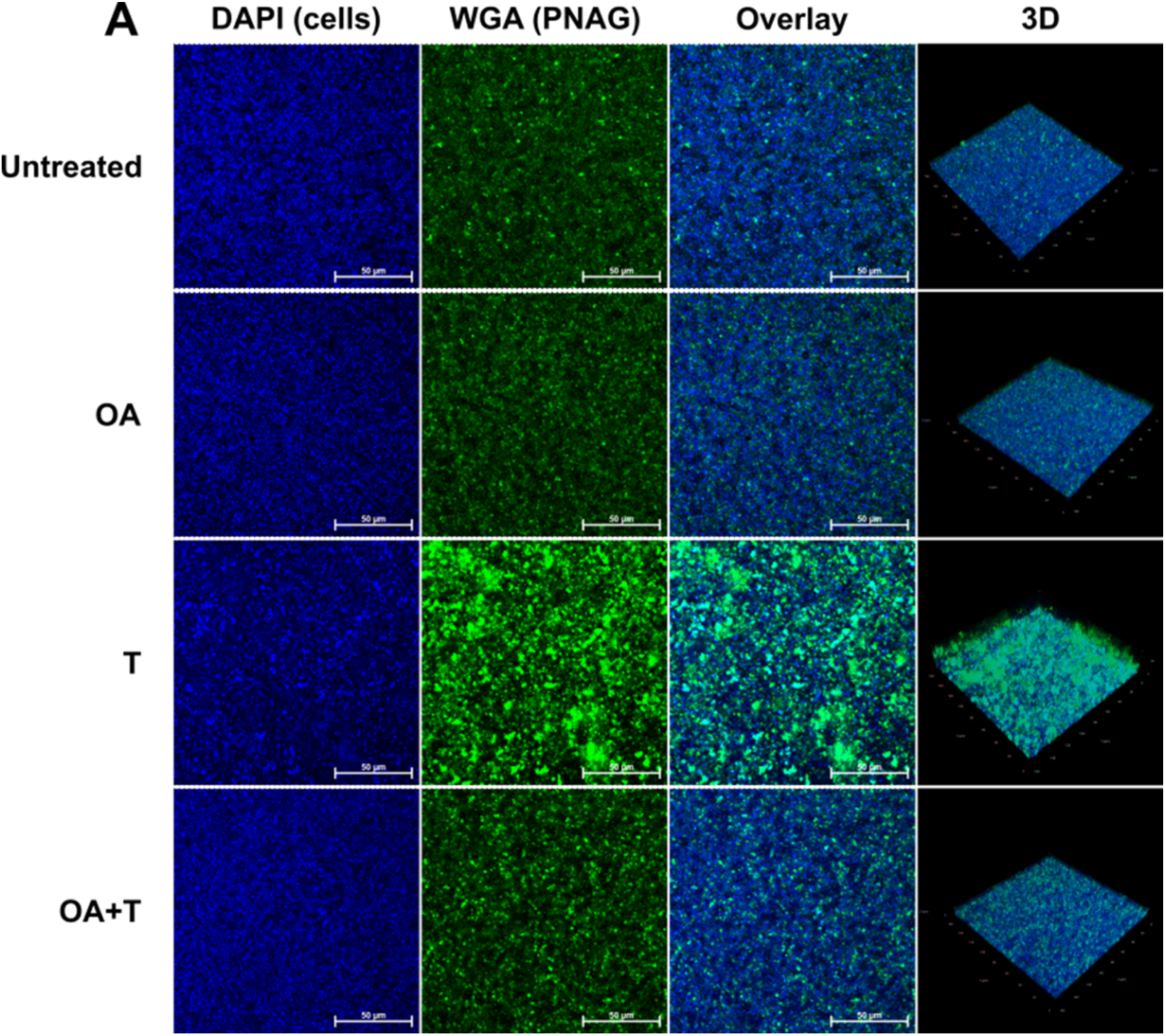
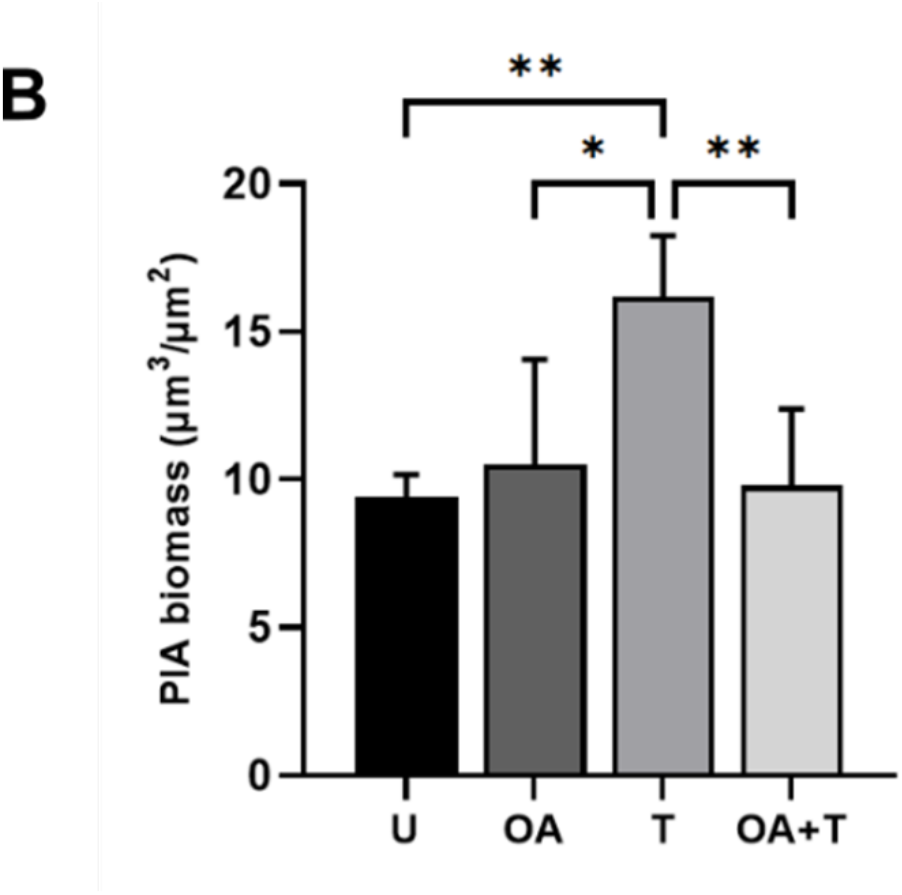
Biofilms grown in the presence of physiologically relevant levels of triclosan produce more matrix polysaccharide, though oleic acid supplementation prevents this. **A)** CLSM was used to assess the effect of triclosan and oleic acid supplementation on the biofilm formation of *S. aureus* HG001 WT biofilms. Conditions include untreated biofilms (U), biofilms supplemented with 500 μM oleic acid (OA), exposed to 500 ng/mL triclosan (T), supplemented with 500 μM oleic acid and exposed to 500 ng/mL triclosan (OA+T). DAPI (blue) was used to visualise cells, fluorescein-conjugated WGA (green) was used to visualise PNAG residues of polysaccharide. For each condition a 2D image of a selected z-plane is shown for DAPI, WGA, and overlay images. A 3D image of each condition is also shown. Images are representative of multiple experiments and were taken using the 40x objective (n=3). **B)** Quantification of polysaccharide biomass was carried out using Comstat2 image analysis software. Error bars represent SD, n=3 * denotes P≤0.05, ** denotes P≤0.01.

### SarA coordinates triclosan-induced polysaccharide synthesis, but not triclosan-induced antibiotic tolerance

Following the findings that physiologically relevant levels of triclosan could induce antibiotic tolerance and alter biofilm formation, the molecular mechanism behind these changes was investigated. The effect of the staphylococcal accessory regulator (Sar) was first explored, as Sar controls the production of PIA in the biofilm matrix of *S. aureus*. Furthermore, PIA can impede the penetration and killing of numerous antibiotics – including vancomycin, ciprofloxacin, and rifampicin (54). Using a *sarA* mutant strain in planktonic kill-curve experiments, *sar* was ruled out from orchestrating triclosan-induced antibiotic tolerance since the triclosan exposed *sarA* mutant, like the WT, exhibited tolerance to vancomycin and ciprofloxacin: ∼3 log fold higher CFUs were seen in *sarA* T+C compared to *sarA* C (Fig 4B) and in *sarA* T+V compared to *sarA* V (Fig 4D). There were no notable differences in the growth of the WT and *sarA* T+C compared to *sarA* across any of the treatments (Fig 4A, 4C).

**Figure 4.**
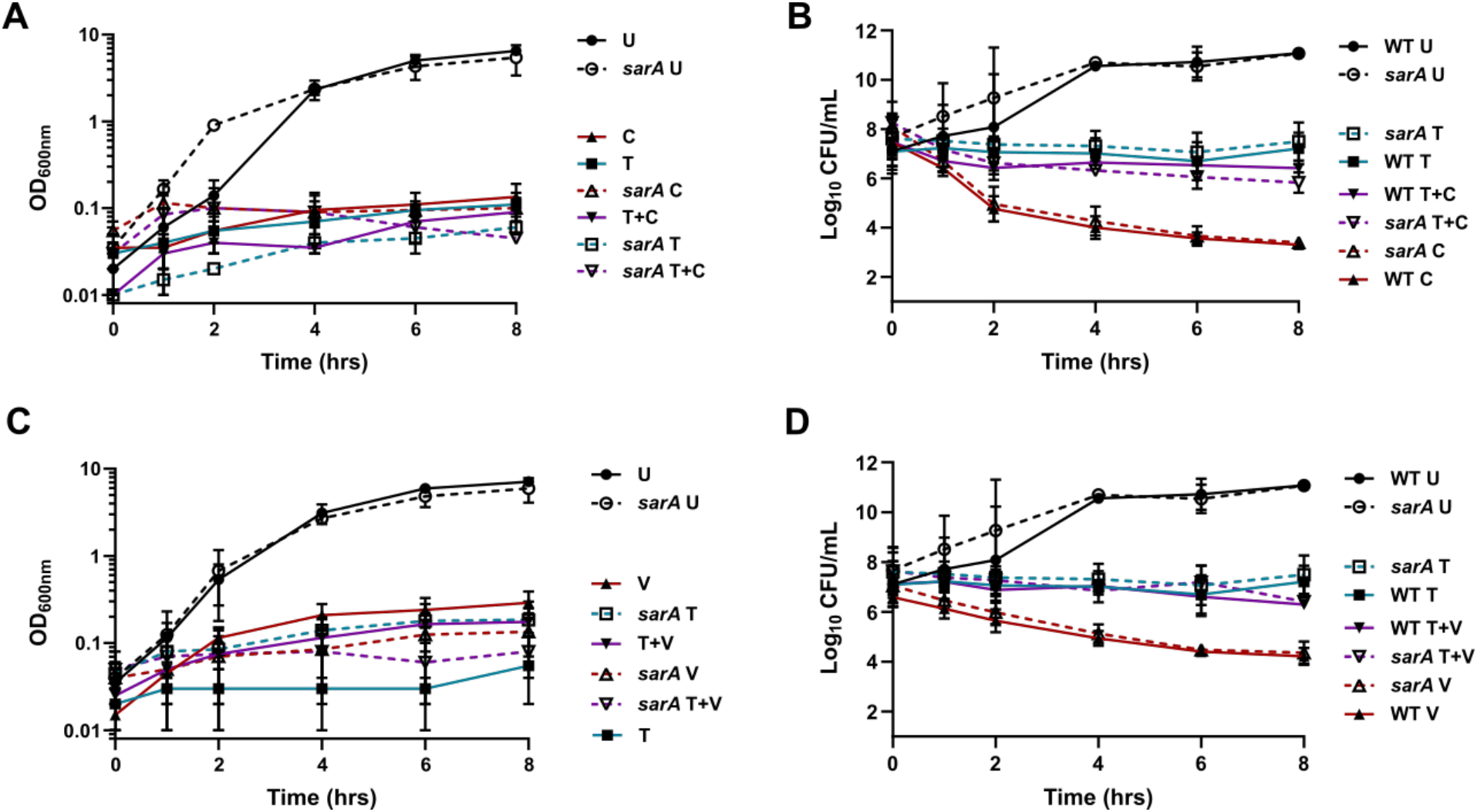
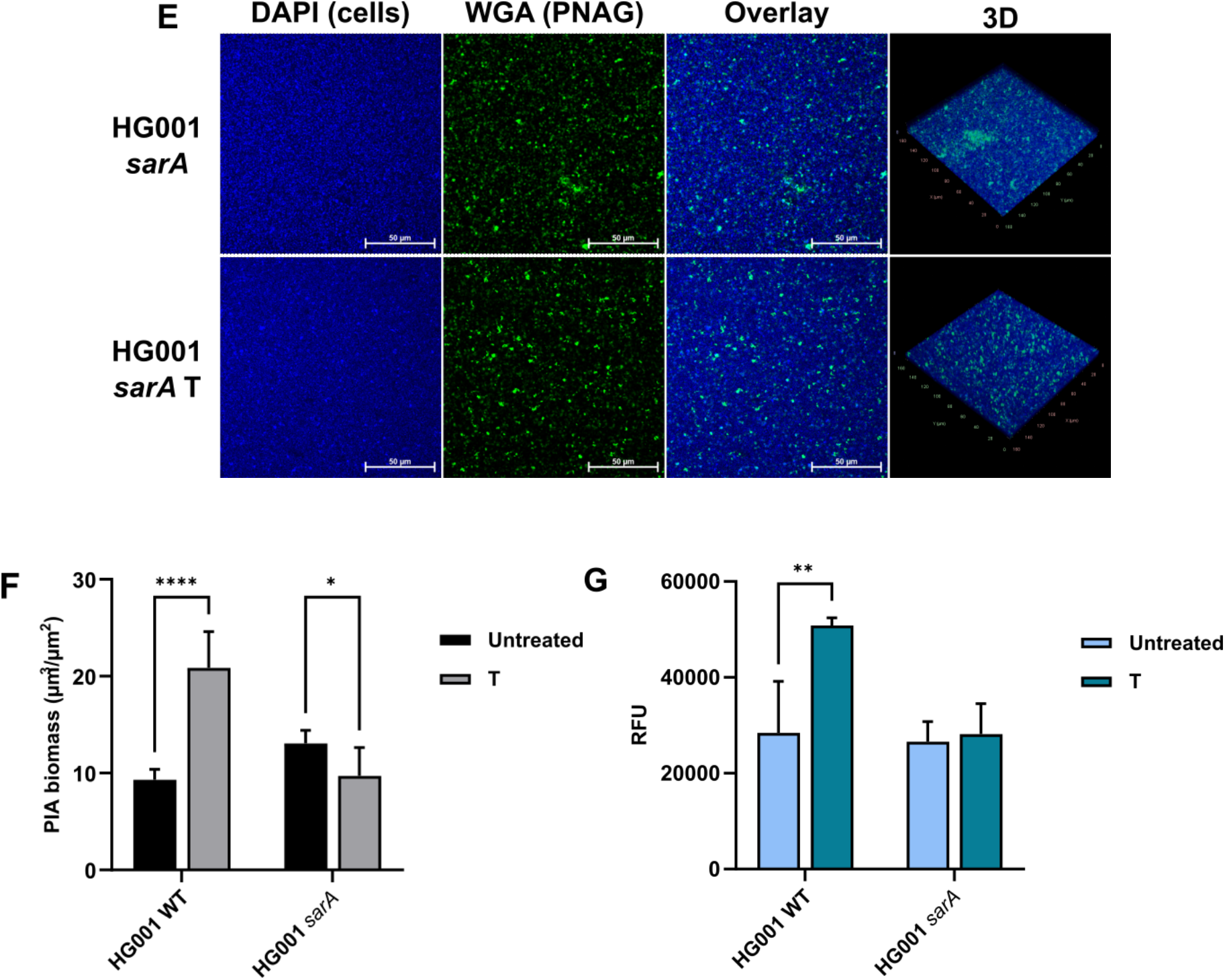
SarA plays no role in triclosan-induced antibiotic tolerance but is responsible for triclosan induced polysaccharide production in *S. aureus* biofilms. *S. aureus* strains HG001 WT and HG001 *sarA* were incubated in BHI in the presence or absence of triclosan for 30 mins at 37°C. Inhibitory concentrations of antibiotic were added and incubation continued. For both strains, untreated (U, *sarA* U), 500 ng/mL triclosan pretreated (T, *sarA* T), 1 μg/mL ciprofloxacin/2 μg/mL vancomycin treated (respectively: C, V, *sarA* C, *sarA* V), 500 ng/mL triclosan treated and 1 μg/mL ciprofloxacin/2 μg/mL vancomycin treated (T+C, T+V, *sarA* T+C, *sarA* T+V) conditions are shown. **A, C)** Growth curve showing optical densities (OD_600_) plotted against time. Error bars display ±SEM, n=3. **B)** Ciprofloxacin and **D)** vancomycin time-kill assays with Log_10_ CFUs plotted against time. Error bars display ±SEM, n=3 (ciprofloxacin, vancomycin). For biofilm experiments, **E)** CLSM was used to assess the effect of triclosan on the biofilm formation of *S. aureus* HG001 *sarA* biofilms. Conditions include untreated biofilms (HG001 *sarA*) and biofilms exposed to 500 ng/mL triclosan (HG001 *sarA* T). DAPI (blue) was used to visualise cells, fluorescein-conjugated WGA (green) was used to visualise PNAG residues of polysaccharide. For each condition a 2D image of a selected z-plane is shown for DAPI, WGA, and overlay images. A 3D image of each condition is also shown. Images are representative of multiple experiments and were taken using the 40x objective (n=3). **F)** Quantification of polysaccharide biomass in HG001 WT and HG001 *sarA* biofilms was carried out using Comstat2 image analysis software. **G)** Quantification of the fluorescence intensity of matrix polysaccharide stained with calcofluor white. Error bars represent SD, n=3. **** denotes P≤0.0001, ** denotes P≤0.01, * denotes P≤0.05.

Whilst SarA did not appear to affect triclosan-induced antibiotic tolerance, imaging *sarA* biofilms using confocal microscopy revealed that triclosan exposure did not result in an increase in the quantity of polysaccharide within the biofilm matrix (Fig 4E), unlike in the WT, in which triclosan exposure resulted in significantly higher proportions of PIA in the biofilm matrix (Fig 4F). Calcofluor white staining of biofilms, in which the binding of the stain to β(1→4) linked D-glucose or derivatives was used to quantify polysaccharide concentrations, confirmed the microscopy results (Fig 4G). The lack of increased matrix polysaccharide of the *sarA* mutant biofilms in the presence of triclosan, suggests triclosan-induced stimulation of SarA results in increased polysaccharide in triclosan exposed biofilms.

### SigB affects neither triclosan-induced polysaccharide synthesis nor triclosan-induced antibiotic tolerance, but may induce a slow growth phenotype in response to triclosan

Since it was found that SarA controlled triclosan-induced polysaccharide production, but not triclosan-induced antibiotic tolerance, the investigation continued with a focus on determining whether SigB played any role. SigB was selected due to its role in coordinating *S. aureus* stress responses (55-57) and its positive regulation of SarA (58). It was therefore hypothesised that activation of SigB by triclosan exposure would stimulate polysaccharide synthesis through SigB mediated upregulation of SarA, whilst simultaneously inducing antibiotic tolerance by another SarA-independent mechanism.

However, like SarA, SigB was found not to play a role in triclosan-induced antibiotic tolerance as triclosan triggered antibiotic tolerance in both a Δ*sigB* mutant and the WT in planktonic kill-curve experiments against ciprofloxacin (Fig 5B) and vancomycin (Fig 5D). In addition, there were no differences in the planktonic growth of the Δ*sigB* mutant compared to the WT in any of the conditions (Fig 5A, 5C). Furthermore, when using *S. aureus* 8325-4 (a strain in which SigB does not function (59)), confocal microscopy (Fig S4A, S4B) and calcofluor white staining (Fig S4C) showed that triclosan exposure still resulted in excess matrix polysaccharide accumulation. This suggests that SigB plays no role in triclosan-induced polysaccharide production, and that the control of triclosan-induced polysaccharide production by SarA is SigB independent.

**Figure 5.**
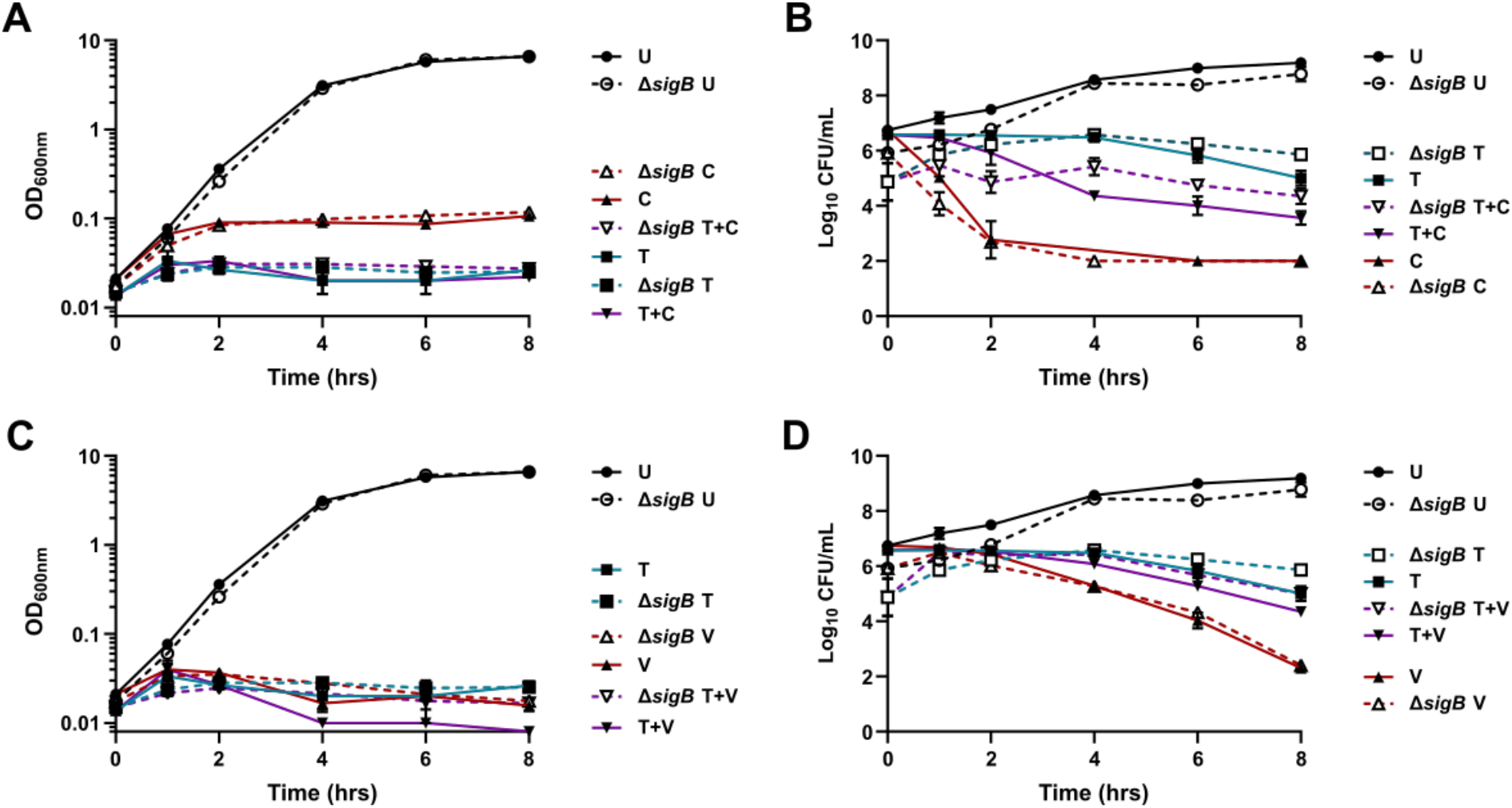
SigB plays no role in triclosan-induced antibiotic tolerance in planktonic *S. aureus* cultures. *S. aureus* strains HG001 WT and HG001 Δ*sigB* were incubated in BHI in the presence or absence of triclosan for 30 mins at 37°C. Inhibitory concentrations of antibiotic were added and incubation continued. For both strains, untreated (U, Δ*sigB* U), 500 ng/mL triclosan pretreated (T, Δ*sigB* T), 1 μg/mL ciprofloxacin/2 μg/mL vancomycin treated (respectively: C, V, Δ*sigB* C, Δ*sigB* V), 500 ng/mL triclosan treated and 1 μg/mL ciprofloxacin/2 μg/mL vancomycin treated (T+C, T+V, Δ*sigB* T+C, Δ*sigB* T+V) conditions are shown. **A, C)** Growth curve showing optical densities (OD_600_) plotted against time. Error bars display ±SEM, n=3. **B)** Ciprofloxacin and **D)** vancomycin time-kill assays with Log_10_ CFUs plotted against time. Error bars display ±SEM, n=3.

### The stringent response is essential for triclosan-induced antibiotic tolerance against multiple, mechanistically different antibiotics

After experiments investigating the action of SarA and SigB in response to triclosan exposure, the mechanism behind triclosan-induced antibiotic tolerance remained elusive. The stringent response has been associated with triclosan-induced antibiotic tolerance in planktonic *E. coli* (36). This study sought to determine whether this was also the case in *S. aureus*. To assess this, an *S. aureus* stringent response (p)ppGpp^0^ mutant incapable of producing (p)ppGpp was subjected to planktonic time-kill assays.

Optical densities displayed the same trends observed in previous experiments, with triclosan pretreatment, antibiotic treatments and triclosan antibiotic combination treatments halting growth. No notable deviations in growth were observed between HG001 WT and HG001 (p)ppGpp^0^ (Fig 6A, 6C). In the WT, triclosan protected *S. aureus* against both bactericidal antibiotics assessed (C, Fig6B; V, Fig 6D). However, the T+C and T+V treatments in the stringent response mutant exhibited 2 log reductions in viability when compared to the WT, indicating triclosan was not able to protect this strain against other antibiotics. This suggests a role for the stringent response in developing antibiotic tolerance following triclosan exposure. Further still, the (p)ppGpp^0^ strain was far more sensitive to killing by triclosan alone, as demonstrated by the 2 log reduction in viability in the (p)ppGpp^0^ T condition vs T (Fig 6B, 6D).

**Figure 6.**
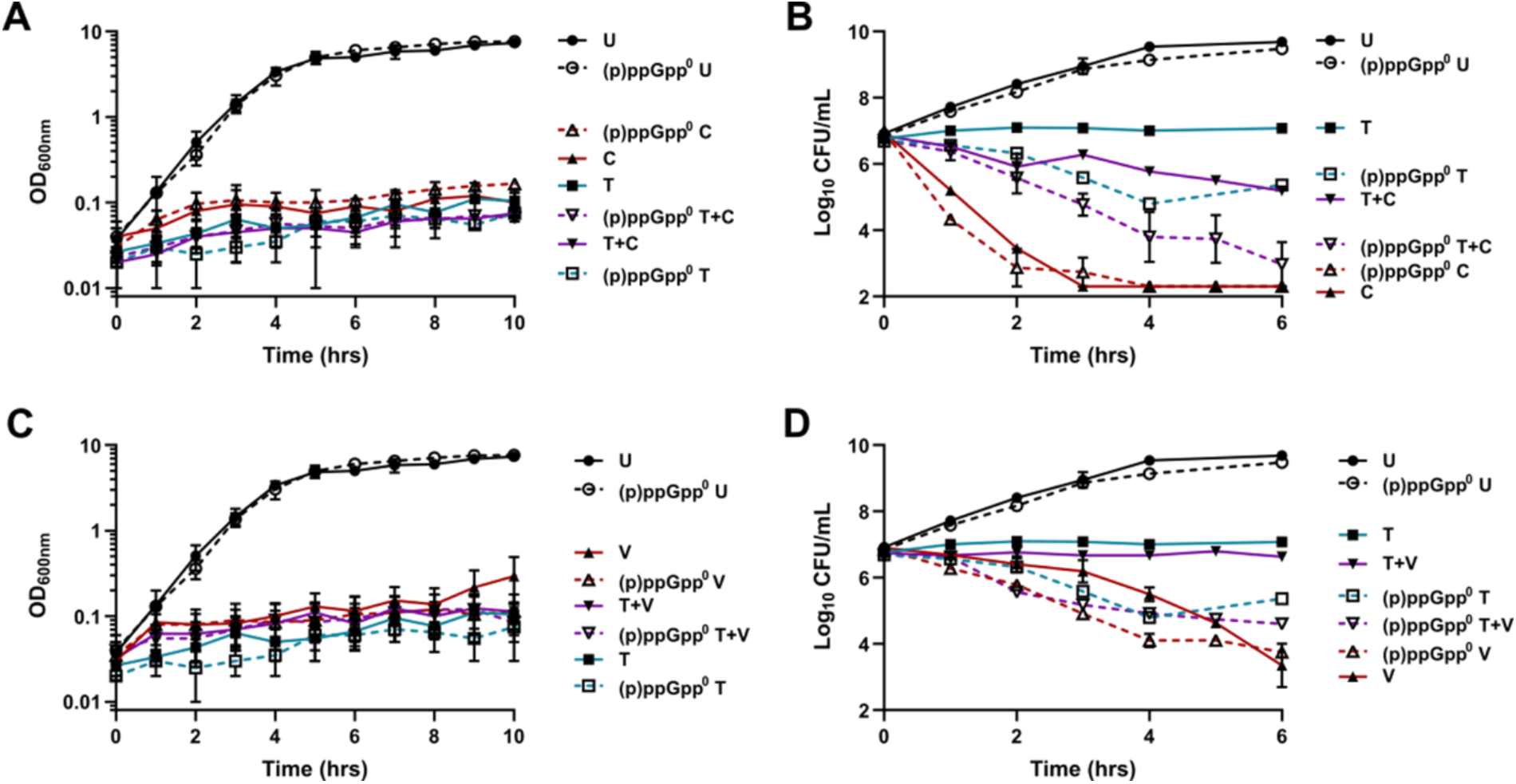
The stringent response regulates triclosan-induced antibiotic tolerance against multiple antibiotics in planktonic *S. aureus*. *S. aureus* strains HG001 WT and HG001 (p)ppGpp^0^ were incubated in BHI in the presence or absence of triclosan for 30 mins at 37°C. Inhibitory concentrations of antibiotic were added and incubation continued. For both strains, untreated (U, (p)ppGpp^0^ U), 500 ng/mL triclosan pretreated (T, (p)ppGpp^0^ T), 1 μg/mL ciprofloxacin/2 μg/mL vancomycin treated (respectively: C, V, (p)ppGpp^0^ C, (p)ppGpp^0^ V), 500 ng/mL triclosan treated and 1 μg/mL ciprofloxacin/2 μg/mL vancomycin treated (T+C, T+V, (p)ppGpp^0^ T+C, (p)ppGpp^0^ T+V) conditions are shown. **A, C)** Growth curve showing optical densities (OD_600_) plotted against time. Error bars display ±SEM, n=2. **B)** Ciprofloxacin and **D)** vancomycin time-kill assays with Log_10_ CFUs plotted against time. Error bars display ±SEM, n=3 (ciprofloxacin, vancomycin).

Biofilm analysis reinforced the importance of the stringent response in triclosan-induced antibiotic tolerance. Live/dead staining shows the (p)ppGpp^0^ strain failing to withstand ciprofloxacin (1× MBEC), vancomycin (1× MBEC), and rifampicin (1× MBEC) treatment despite triclosan exposure throughout biofilm maturation ((p)ppGpp^0^ T+C, Fig S5B; (p)ppGpp^0^ T+V, Fig 7A; (p)ppGpp^0^ T+R, Fig S5C). Comstat2 quantification of live cell biomass shows triclosan significantly protects HG001 WT against ciprofloxacin, vancomycin, and rifampicin (Fig 5B), whilst the (p)ppGpp^0^ T+C, (p)ppGpp^0^ T+V, and (p)ppGpp^0^ T+R biofilms demonstrate levels of viability comparative to WT C, WT V, and WT R. Moreover, the inability of the (p)ppGpp^0^ biofilm to handle stress is highlighted by the reduced viability of the untreated ((p)ppGpp^0^) and triclosan exposed ((p)ppGpp^0^ T), relative the same conditions in the WT (WT, WT T; Fig S5A, 7B).

**Figure 7.**
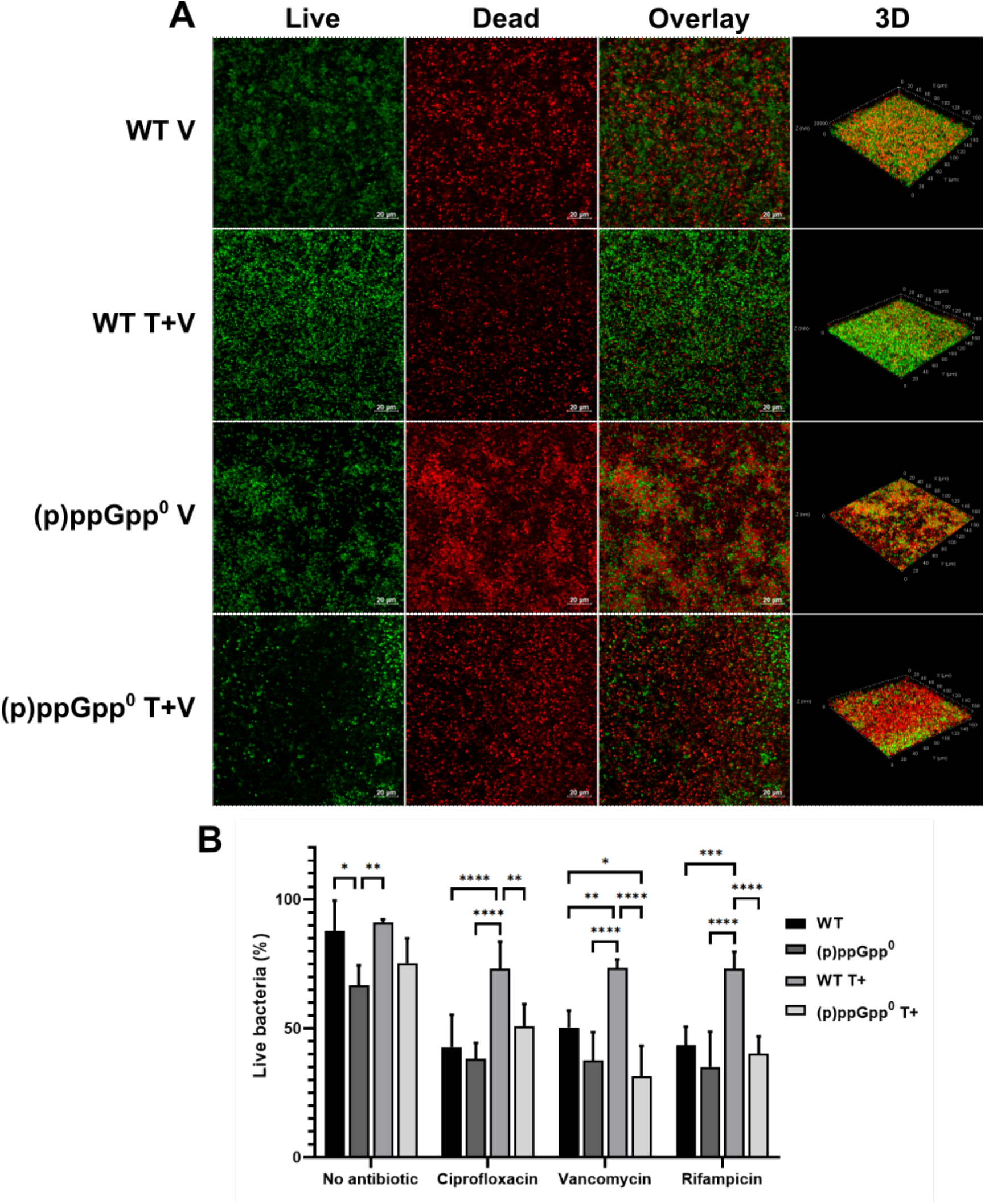
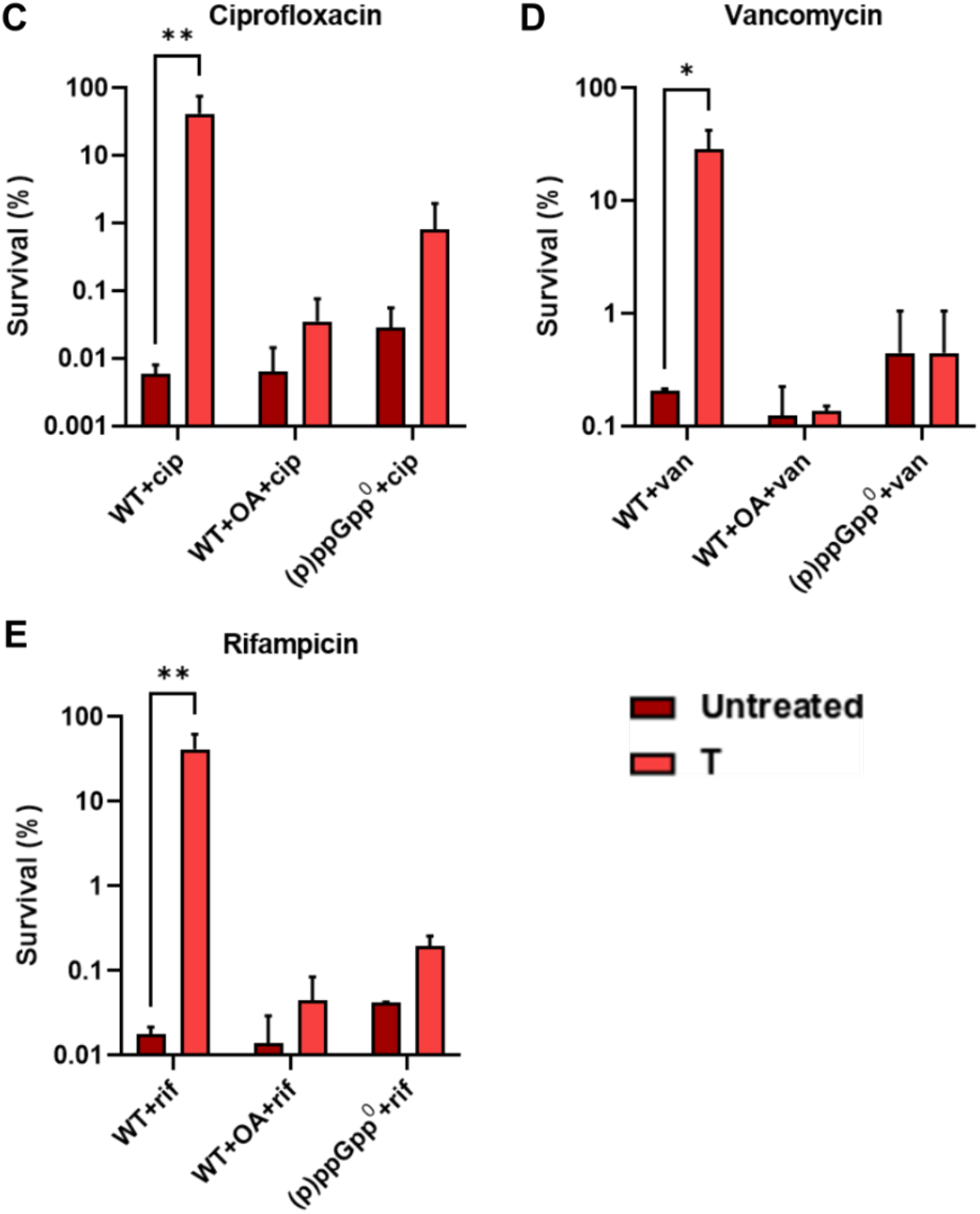
The stringent response plays a role in triclosan-induced antibiotic tolerance against multiple antibiotics in *S. aureus* biofilms. CLSM was used to assess the effect of triclosan on the antibiotic tolerance of *S. aureus* HG001 WT and HG001 (p)ppGpp^0^. Syto 9 (green) was used to visualise live cells, whilst propidium iodide (red) was used to visualise cell death. **A)** Biofilms treated with 2048 μg/mL vancomycin (WT V, (p)ppGpp^0^ V) and 500 ng/mL triclosan exposed biofilms treated with 2048 μg/mL vancomycin (WT T+V, (p)ppGpp^0^ T+V) are shown. For each condition a 2D image of a selected z-plane is shown for live, dead, and overlay images. A 3D image of each condition is also shown. Images are representative of multiple experiments and were taken using the 40x objective (n = 3). **B)** Quantification of the percentage of live cells in biofilms was carried out using Comstat2 image analysis software. Endpoint CFUs of *S. aureus* HG001 (WT), HG001 + 500 μM oleic acid (WT+OA), HG001 (p)ppGpp^0^ ((p)ppGpp^0^) biofilms. Each strain/condition is either untreated or exposed to 500 ng/mL triclosan (T). **C)** Viability of *S. aureus* biofilms following 12 hours of treatment with 4096 μg/mL ciprofloxacin (cip). **D)** Viability of *S. aureus* biofilms following 12 hours of treatment with 2048 μg/mL vancomycin (van). **E)** Viability of *S. aureus* biofilms following 12 hours of treatment with 2048 μg/mL rifampicin (rif). Error bars represent SD, n=3. **** denotes P≤0.0001, *** denotes P≤0.001, ** denotes P≤0.01, * denotes P≤0.05.

Endpoint CFUs of *S. aureus* biofilms also verified that for the HG001 WT, triclosan exposed biofilms were protected against killing by high concentrations of ciprofloxacin (Fig 7C), vancomycin (Fig 7D), and rifampicin (Fig 7E). Additionally, neither oleic acid supplementation nor the use of the (p)ppGpp^0^ mutant resulted in triclosan-induced antibiotic tolerance, further validating the live/dead imaging. In HG001 WT, triclosan was able to induce a 1000-fold increase in tolerance to ciprofloxacin (Fig 7C) and rifampicin (Fig 7E), and a 100-fold increase in vancomycin tolerance (Fig 7D). These findings are even more striking when considering that triclosan pretreated biofilms had been exposed to otherwise lethal concentrations of antibiotics for 12 hours, as opposed to the 3 hours of antibiotic treatment seen in the live/dead biofilm experiments. This could suggest that triclosan exposed *S. aureus* can withstand treatment with extensive levels of antibiotics for prolonged periods of time. Despite these significant changes to antibiotic susceptibility in (p)ppGpp^0^ biofilms, triclosan exposure still resulted in excess polysaccharide being produced in these biofilms (Fig S4A, S4B, S4C), further underlining that triclosan-induced antibiotic tolerance and triclosan-induced polysaccharide production are distinct mechanisms.

## Discussion

This study aimed to characterise the effects of physiologically relevant levels of triclosan on *S. aureus*. Here, we show that 500 ng/mL triclosan, well within the limits of triclosan previously detected in human urine (2.4 - 3,790 ng/mL) (51), can trigger antibiotic tolerance in both planktonic and biofilm culture, and alter biofilm formation. Whereas triclosan induced biofilm formation was dependent on the global regulator SarA, triclosan induced antibiotic tolerance was (p)ppGpp dependent. Triclosan induced biofilm formation as well as antibiotic tolerance were remediated by oleic acid, demonstrating that interruption of fatty acid biosynthesis is the main mode of triclosan action.

Antimicrobial induced biofilm formation has been previously described (37, 40, 60, 61). Here, we shed light on underlying mechanisms by demonstrating that triclosan increases the proportions of polysaccharide present in the biofilm matrix of *S. aureus*. Typically, biofilm formation is stimulated by sub-MIC levels of an antibiotic (60, 61). However, in this study, 500 ng/mL of triclosan, whilst a physiologically relevant concentration of the biocide, is not sub-MIC. This is evidenced by triclosan halting the growth of planktonic cultures and decreasing biofilm cell density. A decrease in cell mass and concurrent stimulation of biofilm matrix production is contrary to many examples of antibiotic induced biofilm formation, but is not a complete anomaly. Skogman and colleagues (2012) found similar results when treating *S. aureus* biofilms with penicillin. They hypothesised that some antimicrobials decrease cell viability whilst increasing production of biofilm matrix components. However, these changes can only be confirmed through parallel measurement of biofilm viability, biomass, and quantifying matrix components, as in this study. Therefore, it may be that more antimicrobials previously associated with stimulating biofilm formation fall into this category, but this is yet to be fully characterised (62). Live/dead staining of triclosan exposed biofilms did not detect any notable increase in the proportions of dead cells. Instead, it appears that triclosan exposed biofilms consist of a reduced population that produces and exports far more PIA, relative to unexposed biofilms.

SigB and Sar are known regulators of *icaADBC* expression, thereby altering the production of PIA synthesis enzymes (63-68). Triclosan induced biofilm formation was independent of SigB. Although SigB acts as a repressor of polysaccharide dependent biofilm formation in *S. aureus* (69), Pant and Eisen (2021) reported that SigB only has an effect on PIA production in particular adverse conditions, such as osmotic stress (68). In contrast, SarA played a key role in triclosan-induced PIA overproduction since a *sarA* mutant was unable to overproduce PIA following triclosan exposure. PIA overproduction could be beneficial for numerous reasons (54), including increased tolerance to mechanical forces and resistance to immunological stresses, such as killing by host antimicrobial peptides and polymorphonuclear leukocytes (70, 71), opsonisation by antibodies and complement (72-74), and phagocytosis by macrophages (70). PIA producing Staphylococci have previously been shown to be less susceptible to killing by some antibiotics, including vancomycin and ciprofloxacin (54, 75, 76). Accordingly, this study hypothesised that triclosan-induced polysaccharide production protected *S. aureus* from killing by antibiotics, and was therefore the cause of triclosan-induced antibiotic tolerance. However, since the *sarA* mutant displayed antibiotic tolerance despite no longer producing excess polysaccharide, there does not appear to be a direct link between polysaccharide production and antibiotic tolerance following triclosan exposure.

Triclosan induced antibiotic tolerance was orchestrated by the stringent response. A (p)ppGpp^0^ strain still overproduces PIA upon triclosan treatment, but no longer benefits from triclosan-induced antibiotic tolerance. Westfall et al. (2019) found that triclosan pretreatment protected planktonic *E. coli* against ampicillin, kanamycin, streptomycin and ciprofloxacin (36) and that triclosan-induced tolerance was mediated by the stringent response. In *S. aureus*, triclosan exposure protected not only planktonic *S. aureus* from ciprofloxacin and vancomycin, but also increased the tolerance of *S. aureus* biofilms against high doses of ciprofloxacin, vancomycin, and rifampicin. (p)ppGpp can specifically block replication, translation and transcription (16, 77). Whether and how triclosan specifically activate (p)ppGpp synthetase remains unclear. The observation that *Bacillus subtilis* fatty acid starvation seems to activate the Rel dependent stringent response (although (p)ppGpp levels remain below the detection limit) could offer a clue (78). The combination of SarA-dependent PIA production to protect against immunological threats, and the stringent response to protect against antimicrobial threats, can lead not only to relapsing infections, but also accelerate the evolution of antibiotic resistance (79).

The triclosan-induced changes to *S. aureus* physiology appear to originate from fatty acid starvation. When growth medium is supplemented with concentrations of oleic consistent with those found in human serum (53), the antibiotic susceptibility of triclosan pretreated *S. aureus* HG001 is restored and biofilm formation unchanged relative to untreated controls. The restoration of antibiotic susceptibility when fatty acid starvation is negated is logical, as fatty acid starvation is one of the numerous nutritional deficiencies capable of instigating the stringent response. However, the link between fatty acid starvation and SarA is less clear, and may suggest the effects of fatty acid starvation are broader than previously thought. The observation that oleic acid supplementation was able to override triclosan-induced affects at all is striking, as the notion that exogenous fatty acids can overcome the effects of fatty acid synthesis inhibitors has been viewed as controversial (80-84). Since the concentration of triclosan used in this experiment was low in comparison to triclosan concentrations in healthcare and household products, it cannot be concluded whether fatty acid supplementation is sufficient to save *S. aureus* from higher concentrations of triclosan. However, the data does suggest serum concentrations of oleic acid (53) would be sufficient to overcome tolerance induced by concentrations of triclosan that have accumulated in the human body (33, 35).

This present study shows that exposure to physiologically relevant levels of triclosan can drive *S. aureus* to trigger multiple, divergent stress responses that alter numerous facets of *S. aureus* physiology. These physiological changes are rooted in the stress caused by triclosan induced fatty acid starvation, before branching off. These diverging responses provide protection against antibiotics, facilitated by the stringent response, and potentially protect from other threats mediated by SarA controlled polysaccharide synthesis.

Skogman and colleagues (2012) advised that the criteria for determining an effective antimicrobial treatment should be based on bacterial viability, biofilm biomass, plus matrix composition. We suggest going further to incorporate potential antibiotic tolerance. If the concentration of triclosan used in this study was to be evaluated based only on biomass and viability, triclosan would be deemed an effective therapy. However, when factoring in the increased matrix production and pleiotropic antibiotic tolerance induced by the biocide, triclosan appears far less alluring. This is further compounded by the finding that triclosan-induced affects occurred at an inhibitory concentration, rather than at sub-MIC levels. Thereby emphasising that accumulated or residual antimicrobial in the human body may cause large scale physiological change to pathogens. This reemphasises the need for stricter control on biocide use globally.

## Acknowledgements

DW was funded via Biological Sciences Research Council, UK Doctoral Training Programme studentship (BB/M008770/1: www.bbsrc.ac.uk) held jointly by the School of Life Sciences and School of Pharmacy, University of Nottingham. KH and JA are partly funded by the National Biofilms Innovation Centre (NBIC) which is an Innovation and Knowledge Centre funded by the BBSRC and InnovateUK (Award Number BB/R012415/1). We thank the School of Life Sciences Imaging Facility (SLIM) for input on image analysis (particularly Robert Markus, Seema Bagia and Tim Self). CW was funded by Deutsche Forschungsgemeinschaft, Schwerpunktprogramm Spp1879 to CW (Project 423246275) and by infrastructural funding from the Deutsche Forschungsgemeinschaft (DFG), Cluster of Excellence EXC 2124 “Controlling Microbes to Fight Infections” (Project 390838134). The project was also supported by the University of Nottingham and the University of Tübingen’s funding as part of the Excellence Strategy of the German Federal and State Governments, in close collaboration with the University of Nottingham.

