## Supplemental figures and tables for "The biocide triclosan induces (p)ppGpp dependent antibiotic tolerance and alters SarA dependent biofilm structures in *Staphylococcus aureus*"

**This PDF file includes:**

Table 1

Figs. 1 to 5

**Table S1. Average minimum inhibitory concentration, minimum bactericidal concentration (MIC) and minimum biofilm eradication concentration (MBEC) of *S. aureus* strains HG001 and 8325-4 against a panel of antibiotics and biocides.**

| Strain | Antimicrobial | MIC (µg/mL) | MBEC (µg/mL) |
|---|---|---|---|
| ***S. aureus* HG001** | Triclosan | 0.1 | - |
| | Ciprofloxacin | 0.25 | 4096 |
| | Vancomycin | 2 | 2048 |
| | Rifampicin | 0.01 | 2048 |
| ***S. aureus* 8325-4** | Triclosan | 0.05 | - |
| | Ciprofloxacin | 0.25 | 4096 |
| | Vancomycin | 2 | 2048 |
| | Rifampicin | 0.01 | 2048 |

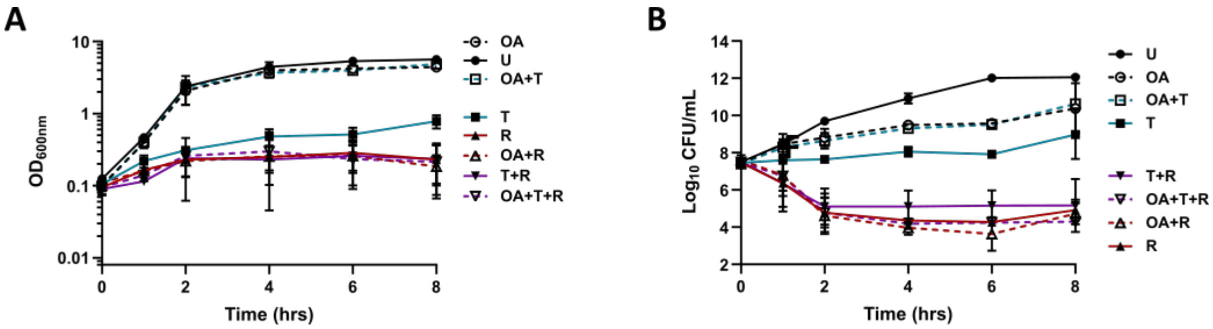


**Supplementary figure 1. Triclosan is unable to induce notable tolerance to bacteriostatic levels of rifampicin in planktonic *S. aureus* cultures.** *S. aureus* strain HG001 was incubated in BHI in the presence or absence of triclosan, with or without 500 µM oleic acid solubilised in 0.1% Brij 58, for 30 mins at 37°C. Inhibitory concentrations of antibiotics were added and incubation continued. Untreated (U, OA), 500 ng/mL triclosan pretreated (T, OA+T), 40 ng/mL rifampicin treated (R, OA+R), 500 ng/mL triclosan and 40 ng/mL rifampicin treated (T+R, OA+T+R) conditions are shown. Growth curve with rifampicin **(A)** showing optical densities ($OD_{600}$) plotted against time. Error bars display ±SD, n=3. **B)** Rifampicin time-kill assays with $Log_{10}$ CFUs plotted against time. Error bars display ±SD, n=3.

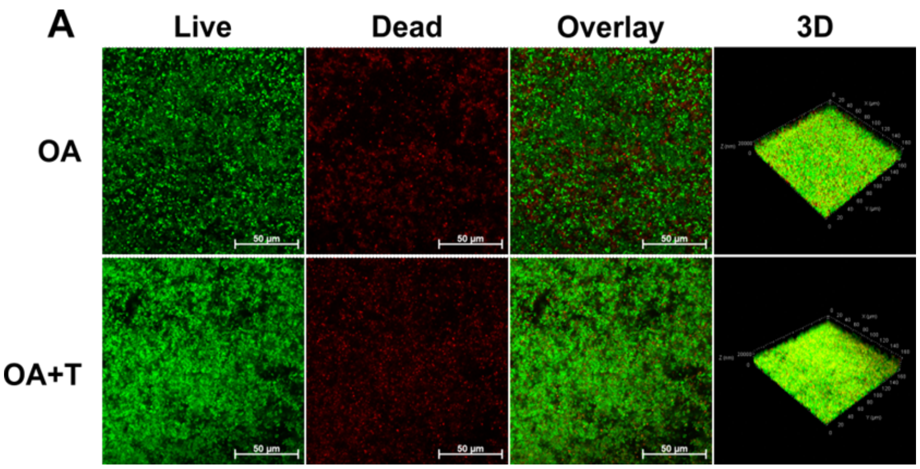

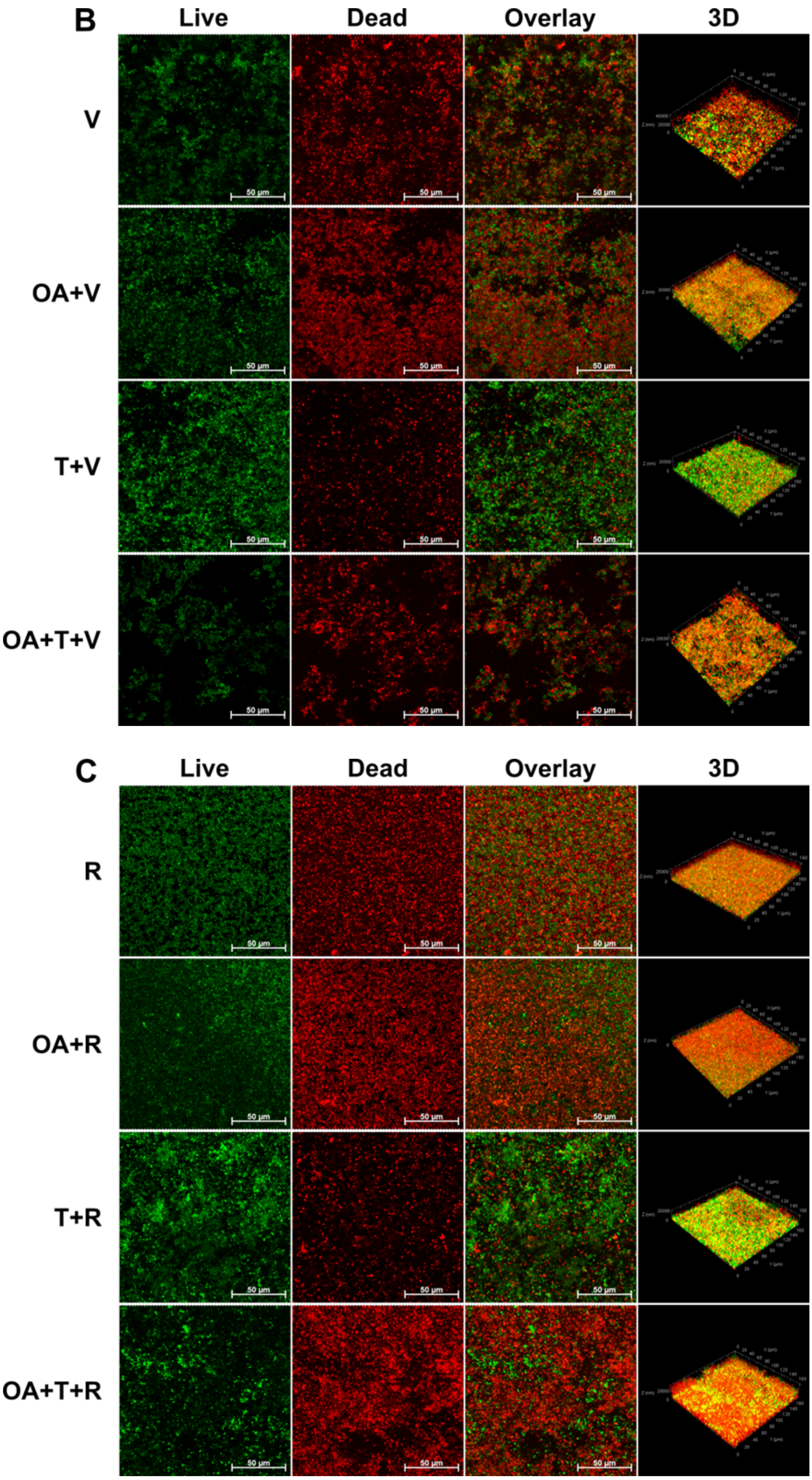


**Supplementary figure 2. Triclosan exposure increases biofilm tolerance to vancomycin and rifampicin, but susceptibility can be restored with oleic acid supplementation**. CLSM was used to assess the effect of triclosan on the antibiotic tolerance of *S. aureus* HG001 WT. Syto 9 (green) was used to visualise live cells, whilst propidium iodide (red) was used to visualise cell death. **A)** Biofilms grown in the presence of 500 µM oleic acid, untreated (OA) and 500 ng/mL triclosan exposed (T, OA+T) are shown. **B)** Biofilms grown in the presence or absence of 500 µM oleic acid and treated with 2048 µg/mL vancomycin (V, OA+V) and 500 ng/mL triclosan exposed biofilms treated with 2048 µg/mL vancomycin (T+V, OA+T+V) are shown. **C)** Biofilms grown in the presence or absence of 500 µM oleic acid and treated with 2048 µg/mL rifampicin (R, OA+R) and 500 ng/mL triclosan exposed biofilms treated with 2048 µg/mL rifampicin (T+R, OA+T+R) are shown.  For each condition a 2D image of a selected z-plane is shown for live, dead, and overlay images. A 3D image of each condition is also shown. Images are representative of multiple experiments and were taken using the 40x objective (n = 3).

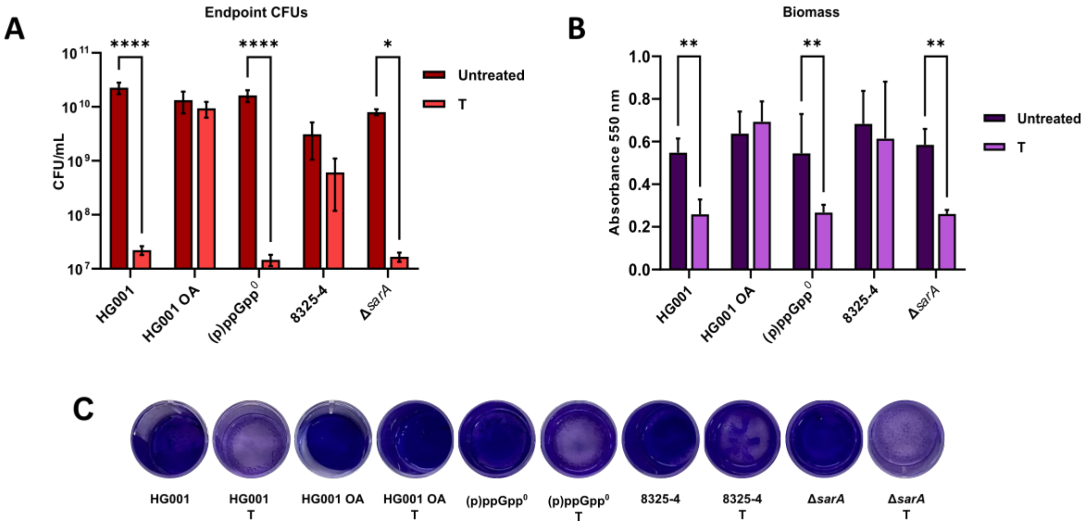


**Supplementary figure 3. SigB appears to control the switch to slow growth induced by physiologically relevant levels of triclosan. A)** Endpoint CFU/mL of biofilms grown for 48 h. **B)** Biomass of biofilms measured by spectral absorption following crystal violet staining. **C)** Photographs of crystal violet stained biofilms. Error bars represent SD, n=3 **** denotes P≤0.0001, ** denotes P≤0.01, * denotes P≤0.05.

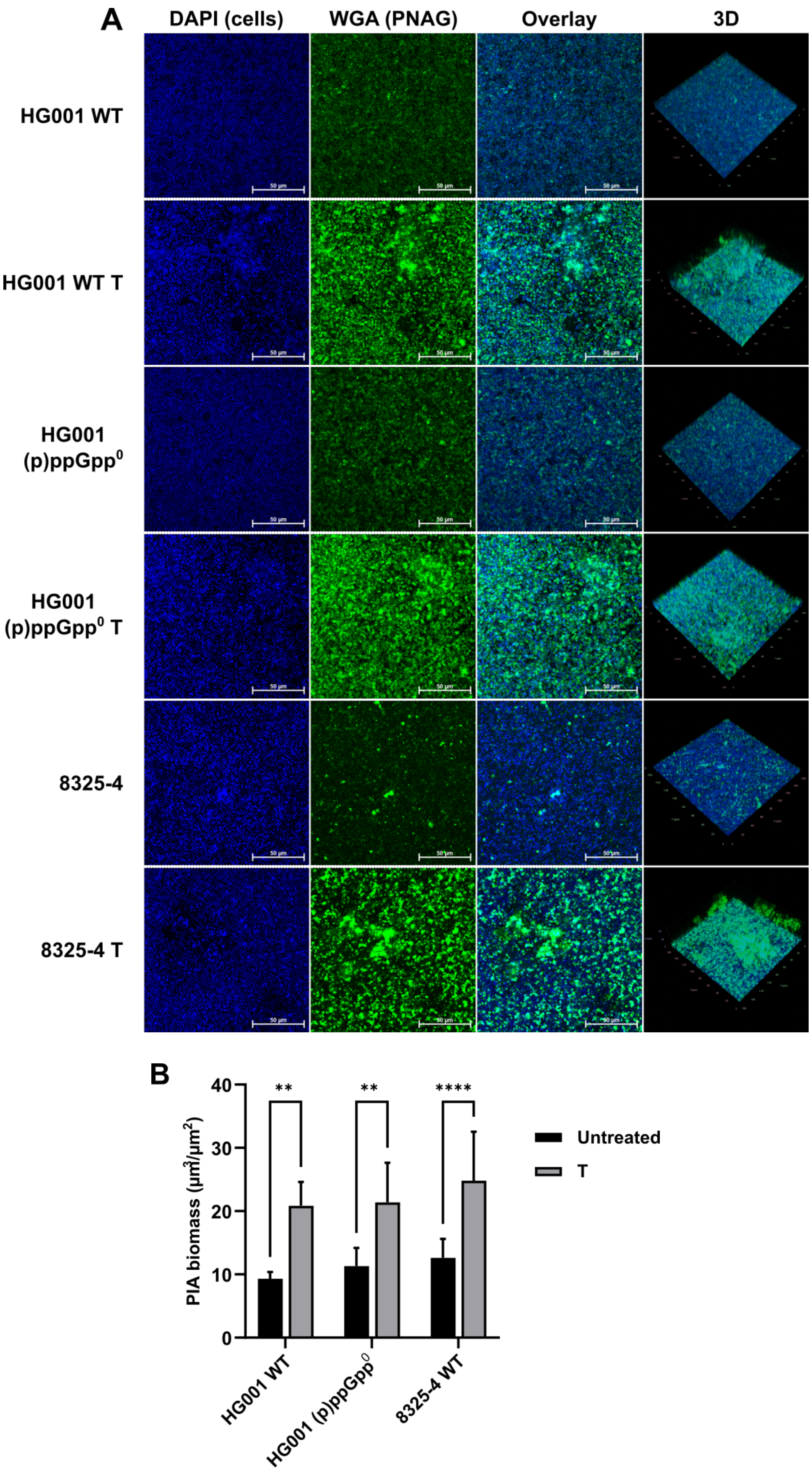
A
DAPI (cells)
WGA (PNAG)
Overlay
3D
HG001 WT
HG001 WT T
HG001 (p)ppGpp0
HG001 (p)ppGpp0 T
8325-4
8325-4 T
50 µm
B
PIA biomass (µm3/µm2)
40
30
20
10
0
**
**
****
Untreated
T
HG001 WT
HG001 (p)ppGpp0
8325-4 WT

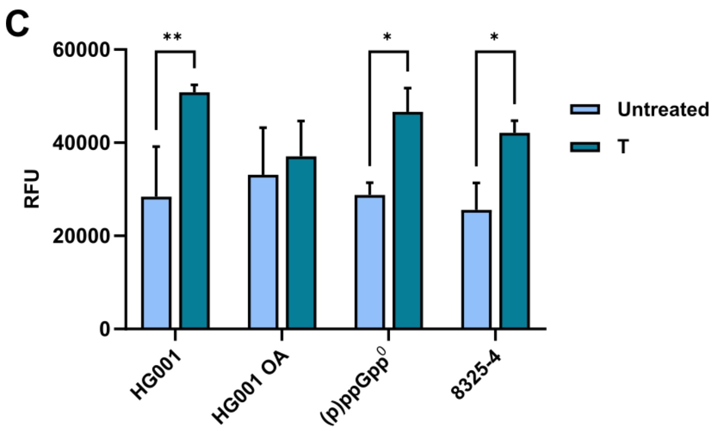


**Supplementary figure 4. SigB and the stringent response have no effect on triclosan-induced polysaccharide production. A)** CLSM was used to assess the effect of triclosan on the biofilm formation of *S. aureus* HG001 WT, HG001 (p)ppGpp$^0$ and 8325-4 WT biofilms. Conditions include untreated biofilms (HG001 WT, HG001 (p)ppGpp$^0$, 8325-4) and biofilms exposed to 500 ng/mL triclosan (HG001 WT T, HG001 (p)ppGpp$^0$ T, 8325-4 T). DAPI (blue) was used to visualise cells, fluorescein-conjugated WGA (green) was used to visualise PNAG residues of polysaccharide. For each condition a 2D image of a selected z-plane is shown for DAPI, WGA, and overlay images. A 3D image of each condition is also shown. Images are representative of multiple experiments and were taken using the 40x objective (n=3). **B)** Quantification of polysaccharide biomass in HG001 WT, HG001 (p)ppGpp$^0$, 8325-4 biofilms was carried out using Comstat2 image analysis software. **C)** Quantification of the fluorescence intensity of matrix polysaccharide stained with calcofluor white. Error bars represent SD, n=3. **** denotes P≤0.0001, ** denotes P≤0.01, * denotes P≤0.05.

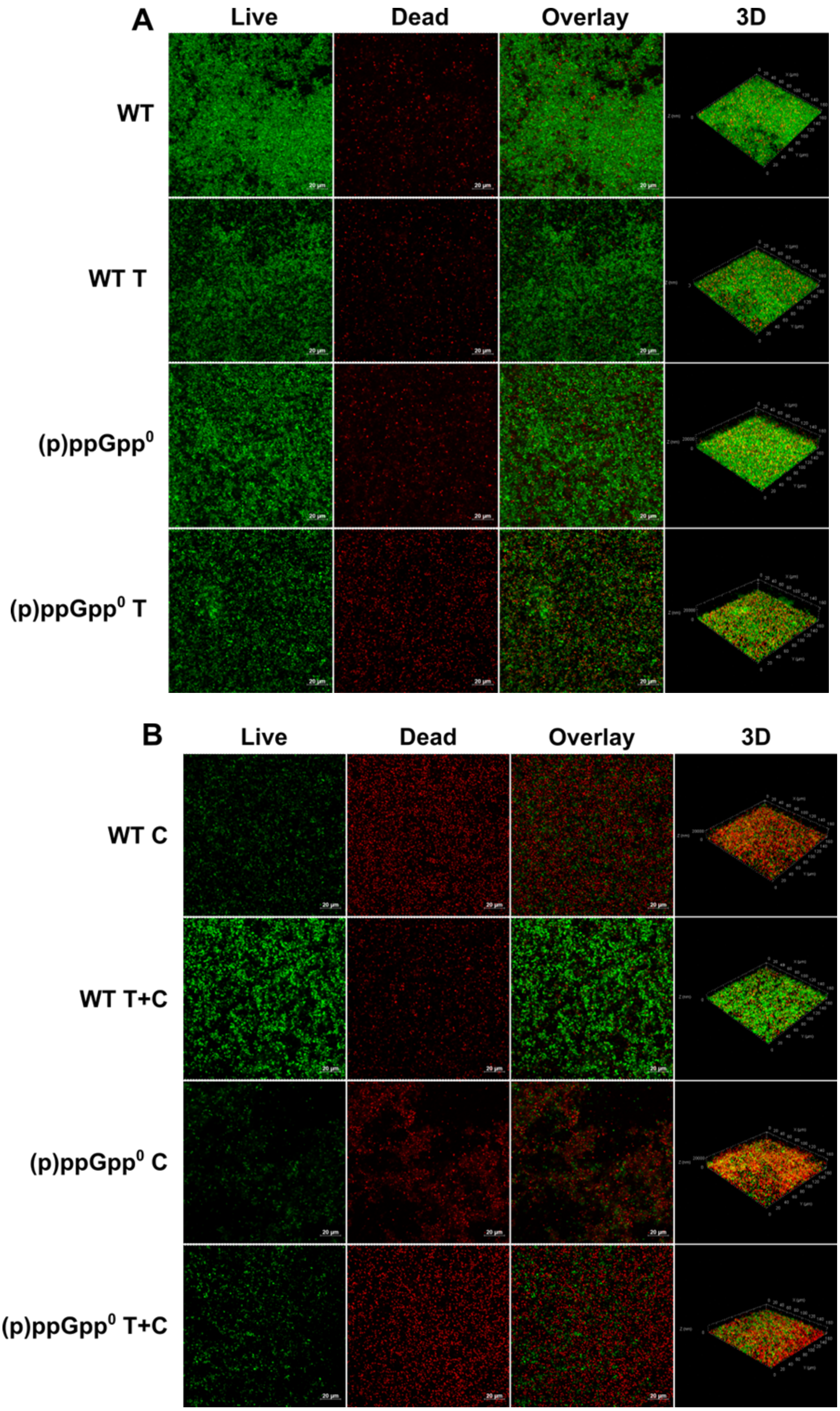
A
Live
Dead
Overlay
3D
WT
WT T
(p)ppGpp⁰
(p)ppGpp⁰ T
B
Live
Dead
Overlay
3D
WT C
WT T+C
(p)ppGpp⁰ C
(p)ppGpp⁰ T+C

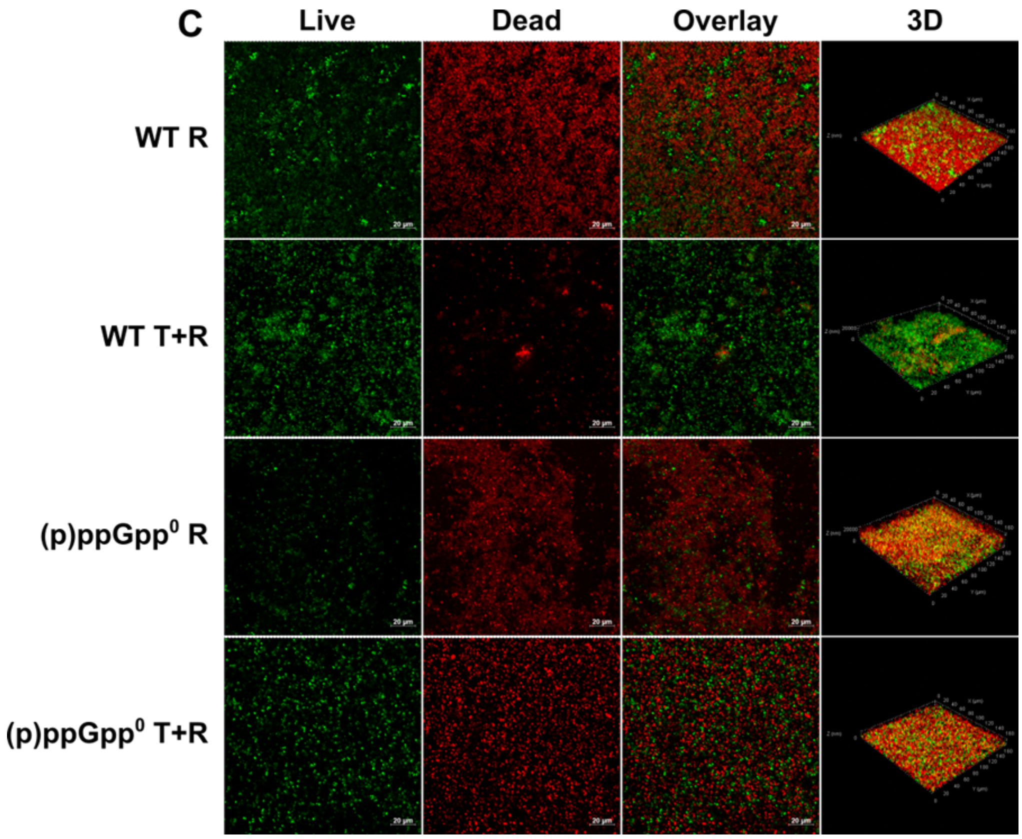


**Supplementary figure 5. Triclosan cannot protect biofilms of the stringent response mutant against killing by ciprofloxacin or rifampicin**. CLSM was used to assess the effect of triclosan on the antibiotic tolerance of *S. aureus* HG001 WT and HG001 (p)ppGpp$^0$. Syto 9 (green) was used to visualise live cells, whilst propidium iodide (red) was used to visualise cell death. **A)** Untreated (WT, (p)ppGpp$^0$) and 500 ng/mL triclosan exposed (WT T, (p)ppGpp$^0$ T) are shown. **B)** Biofilms treated with 4096 μg/mL ciprofloxacin ( WT C, (p)ppGpp$^0$ C) and 500 ng/mL triclosan exposed biofilms treated with 4096 μg/mL ciprofloxacin (WT T+C, (p)ppGpp$^0$ T+C) are shown. **C)** Biofilms treated with 2048 μg/mL rifampicin (WT R, (p)ppGpp$^0$ R) and 500 ng/mL triclosan exposed biofilms treated with 2048 μg/mL rifampicin (WT T+R, (p)ppGpp$^0$ T+R) are shown. For each condition a 2D image of a selected z-plane is shown for live, dead, and overlay images. A 3D image of each condition is also shown. Images are representative of multiple experiments and were taken using the 40x objective (n = 3).
